# Concentration dependent CsrA regulation of the *uxuB* transcript leads to development of post-transcriptional BANDPASS Filter

**DOI:** 10.1101/2024.09.28.615595

**Authors:** Alejandra M. Rojano-Nisimura, Trevor R. Simmons, Alexandra J. Lukasiewicz, Ryan Buchser, Josie S. Ruzek, Jacqueline L. Avila, Lydia M. Contreras

**Author notes:** Corresponding Author: Lydia M. Contreras. These authors contributed equally to this work and are considered co-first authors (Alejandra M. Rojano-Nisimura, Trevor R. Simmons).

## Abstract

Post-transcriptional control systems offer new avenues to design synthetic circuits that offer reduced burden and less synthetic regulatory components compared to transcriptionally based tools. Herein, we repurpose a newly identified post-transcriptional interaction between the *uxuB* leader sequence and the *E. coli* CsrA regulatory protein to design a biological post-transcriptional BANDPASS filter. In this work, we characterize *the uxuB* mRNA as heterogenous target of the Carbon Storage Regulatory A (CsrA) protein, where the protein can both activate and repress *uxuB* activity depending on its intracellular concentration. We leverage this interaction to implement a novel strategy of regulation within the 5’UTR of an mRNA. Specifically, we report a hierarchical binding strategy that may be leveraged by CsrA within *uxuB* to result in a dose-dependent response in regulatory outcomes. In our semi-synthetic circuit, the *uxuB* mRNA leader sequence is used as a scaffold that is fused to a gene of interest, which allows the circuit to transition between ON/OFF states based on a range of free native concentration of the CsrA regulatory protein. Notably, this system exerts regulation comparable to previously developed transcriptional BANDPASS filters while reducing the number of circuit components and can be used in concert with additional controlled circuits to achieve complex multi-signal control. We anticipate that future characterization of native regulatory RNA-protein systems will allow for development of more complex RNP-based circuits for synthetic biology applications.

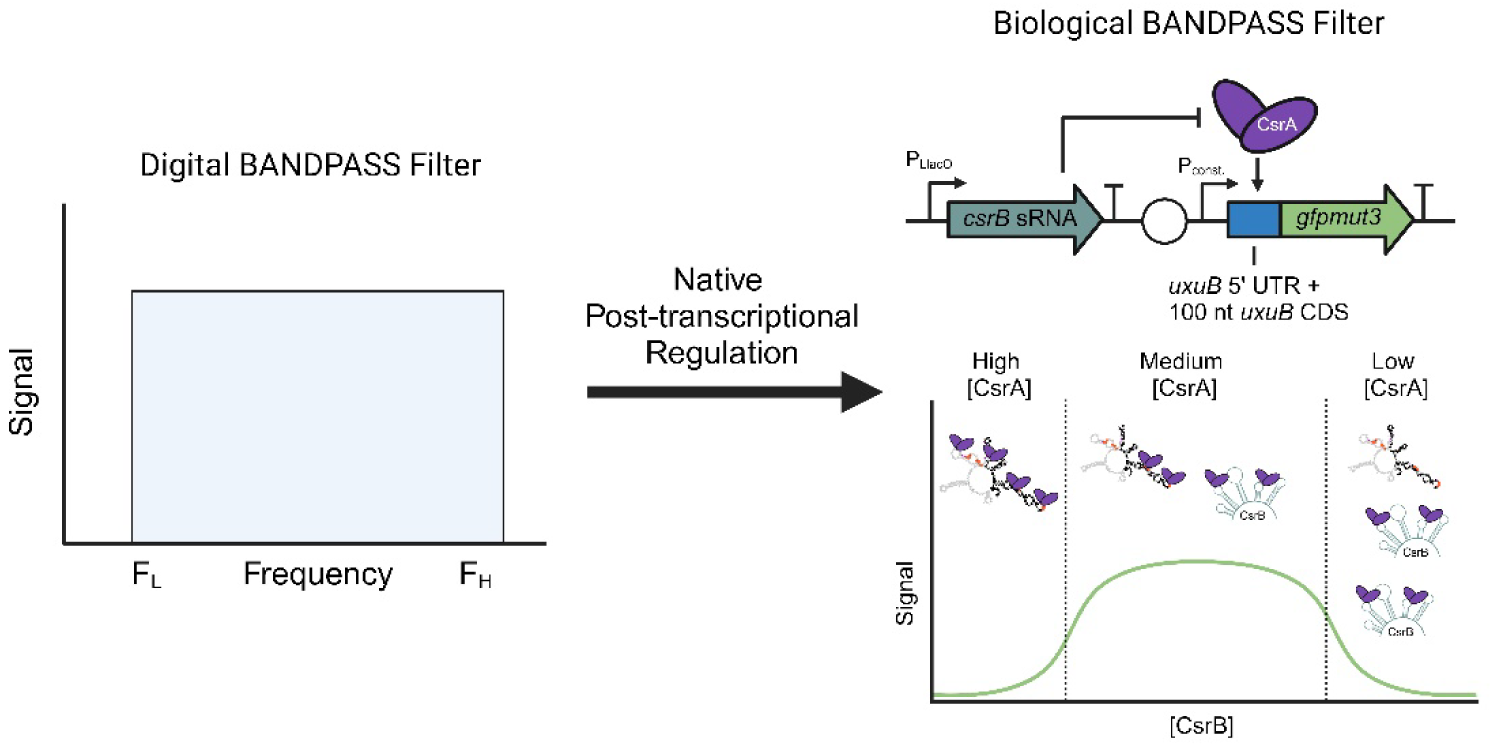

## Introduction

Constructing biological control systems that can process multiple input signals and actuate dynamic genetic outputs (e.g., oscillation, memory, pulsing, etc.) has been a long-standing goal in synthetic biology^1–3^. To achieve these goals, two primary approaches have been utilized: bottom-up and top-down design. Bottom-up designs develop *de novo* systems via synthetic components, while top-down designs repurpose native biological parts and networks to achieve user-defined outcomes^4^. Bottom-up approaches can offer precise tuning between synthetic components, and currently perform effectively in *in vitro* and cell-free settings^5–7^.

Top-down approaches can be more effective *in vivo* due to the benefits conferred by native systems, such as beneficial host environments and the natively evolved function of the genetic components^8^. Previous top-down efforts have primarily utilized transcription-based regulation to create diverse toolkits that can be employed into complex, tunable circuits^9–12^. However, as these circuits increase in complexity, unwanted crosstalk or improper coordination of parts can occur, leading to a reduction in circuit performance or complete system failure^13,14^. Additionally, increasing the complexity and number of circuit parts also results in overburden due to cellular resource limitations and trade-offs in fold change^15,16^.

To avoid crosstalk and burden, native systems can serve as a scaffold to integrate with the host’s native regulatory networks^17,18^. One avenue within this approach is to utilize RNA-based or ribonucleoprotein (RNP)-based modules given their overall reduced cellular burden^19,20^ and the multiple nodes for tuning expression at the post-transcriptional level^21–25^. The Carbon Storage Regulatory (Csr) Network, also known as the Regulatory of Secondary Metabolites (Rsm) Network, offers a varied repertoire of regulatory mechanisms that are natively mediated through RNA-Protein interactions^26,27^. The key regulator of the Csr Network is the Carbon Storage Regulatory A (CsrA) protein, which is a 61 amino acid RNA-binding protein (RBP) that regulates hundreds of RNAs a target-specific manner^28,29^. Canonically, CsrA binds mRNA transcripts within the 5’ untranslated region (5’ UTR) through the first 100 nucleotides (nts) of coding sequence. This interaction typically results in translational repression by occluding the Shine Dalgarno sequence and preventing ribosome binding^30^. It primarily binds to an A(N)GGA motif, where N is any nucleotide, within the stem loop of the 5’ UTR, moving forward we will refer to the A(N)GGA motif simply as the GGA motif. CsrA also can enhance gene expression by altering the secondary structure of the leader sequence to promote ribosome loading as well as by stabilizing RNA to prevent degradation by RNases^31^. CsrA is primarily regulated by two small RNAs (sRNAs), CsrB and CsrC, which sequester up to nine and five copies of the CsrA protein^32,33^. In addition, other CsrA sponges have also been identified^26,34^. Lastly, CsrB and CsrC are regulated by the protein CsrD, which drives degradation via RNase E^35^.

Previously, the Csr network has been implicated in bioprocessing. In McKee et al. 2012, the authors found that overexpressing the CsrB sRNA increased titers of fatty acids and isoprenoids^36^. Moreover, interactions within the Csr network have been shown to be tunable. For instance, Leistra and colleagues rationally engineered a library of CsrB mutants to tune the affinity between CsrA and CsrB for wider synthetic biology applications^37^. Most recently, we designed a collection of Csr-regulated control schemes that leverage native CsrA expression to create post-transcriptionally regulated circuits, such as two-input Boolean logic gates and a genetic pulse^27^. An important benefit of these Csr-based circuits is that they can be easily modified in a predictable fashion that is directly correlated with the native molecular interactions between their basic components (e.g., CsrA-mRNA or CsrA-CsrB interactions).

Previous biophysical modeling of CsrA on predicted mRNA targets identified a subset of 22 out of 107 modeled mRNAs as “heterogeneous targets”^38^, whereby it was predicted that there were at least two CsrA-mRNA bound conformations equally likely to occur but resulting in differential regulatory outcomes (*i.e*., translational activation, or translational repression). These results align with previous work, in which genes that affect cell morphology under stress were found to be differentially regulated by CsrA across growth phases in *Vibrio cholera*^39^. Similar growth-dependent effects of CsrA regulation have also been observed in *Salmonella enterica*^28^. However, no mechanism of differential regulation by CsrA has been identified in these or other previously characterized targets.

In this work, we use a combination of *in vitro* biochemical methods and *in vivo* assays to characterize the interaction between CsrA and the *uxuB* mRNA transcript, which was predicted to be differentially regulated by CsrA based on the specific binding site combinations used^38^. Importantly, we show that CsrA can activate expression of *uxuB* in a dose-dependent manner. We identify two independent binding interfaces, located in the 5’ UTR and in the first hundred nucleotides of coding sequence (CDS), that are responsible for two regulatory modes based on intracellular CsrA availability. *In vitro* studies further confirm that CsrA can bind independently and simultaneously to these two interfaces and provide insights regarding the hierarchy of these binding sites. This represents, to our knowledge, the first example of bi-directional regulation of an mRNA transcript by the CsrA regulatory protein.

The unique ability of CsrA to tune the regulatory outcome of *uxuB* in a concentration-dependent manner makes this system well suited for novel post-transcriptional genetic control schemes. Here, we demonstrate that the *uxuB*-CsrA interaction can be easily adapted to create a tunable BANDPASS filter, which is defined as a circuit that achieves system activation only within a specific range of input signal. This BANDPASS filter responds comparably to previous transcriptionally regulated biological BANDPASS filters and also can be deployed in parallel with other Csr-controlled circuits to achieve complex regulatory outcomes. Additionally, this BANDPASS filter reduces the number of required components by a factor of two, as we leverage the native dose-dependency between CsrA and the *uxuB* transcript. To our knowledge, this is the first demonstration of a post-transcriptional BANDPASS filter in a bacterial system. Through our characterization the *uxuB* transcript and its regulation by CsrA, we demonstrate that the discovery and mechanistic understanding of RNA-Protein interactions offer new opportunities for co-opting native machinery to synthesize complex and efficient biological circuits.

## Results and Discussion

### Screening putative heterogenous mRNA targets of CsrA identifies *uxuB* as a unique regulatory target

In previous work, we utilized a computational biophysical model developed for CsrA to identify 22 putative mRNAs targets that had the potential to be heterogeneously regulated by CsrA^38^. Heterogonous regulation specifically refers to a prediction in which CsrA directly binds an mRNA target in different conformations, and each bound confirmation results in different regulatory outcomes (i.e., repression, activation, or no regulation). Heterogeneous targets are interesting candidates for characterization, as they represent single RNA scaffolds that hold the potential to achieve complex biological computation. Based on the molecular features of these 22 mRNAs (**Supplementary Table S3**), we selected 14 mRNA transcripts to further investigate direct interactions with CsrA *in vivo.* To confirm direct mRNA-CsrA interactions in vivo, we used a three-component fluorescence complementation assay (TriFC) previously developed in our lab (**Figure 1a, b**)^40,41^. In short, the TriFC assay utilizes a split YFP protein to determine CsrA-RNA interactions. CsrA is fused to the N-terminal YFP domain (NYFP), and the MS2 Coat Protein is fused to the C-terminal YFP domain (CYFP). Additionally, the RNA sequence of interest is fused to the 5’ end of the 4x MS2 Binding hairpin. If CsrA binds the RNA of interest, the NYFP and CYFP domains can interact and reconstitute to drive YFP fluorescence. If no RNA-CsrA interaction occurs, fluorescence via complementation in minimal. Through this assay we identified 8 novel direct CsrA targets (*entC, yqjE, rodZ, cmk, dps, ybaL, patA and uxuB*) (**Figure 1c**). We then confirmed CsrA binding using *in vitro* electrophoretic mobility shift assays (**Supplementary Figure S1**) for all targets except *dps*. This suggests there are potentially additional cellular factors needed to mediate the CsrA-*dps* interaction. Additionally, complex formation between CsrA and *patA* was only detected *in vitro*. This could be due to *patA* secondary structure features which can hinder YFP complementation.

**Figure 1.**
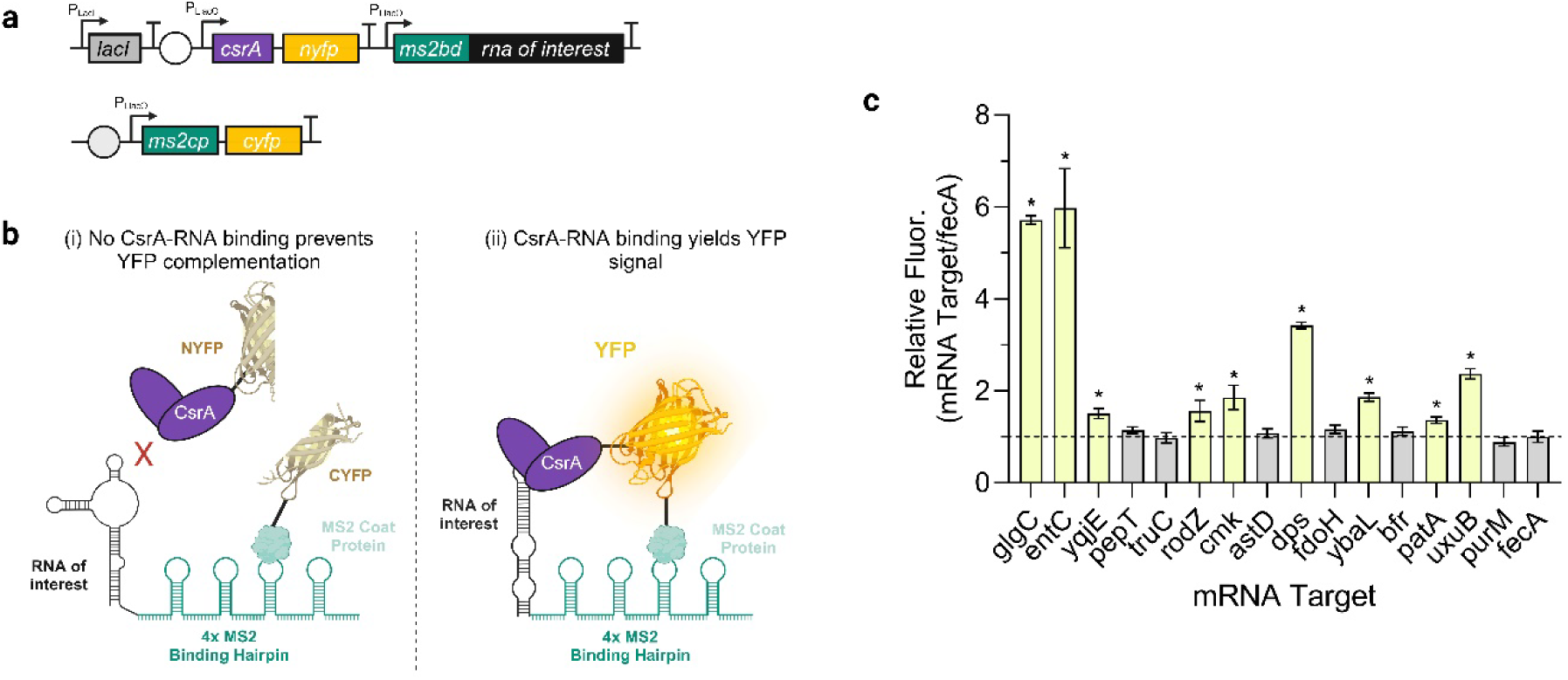
CsrA directly interacts with novel mRNA targets *in vivo*. a) Diagram of the two-plasmid system for *in vivo* three-component complementation assay (TriFC) (adapted from **Leistra and Mihailovic *et al.* 2018**). b) Three-component fluorescence complementation system for detecting direct CsrA-mRNA binding consists of the leader sequence of the mRNA of interest fused to the MS2 binding domain and the *rrnB* terminator, a MS2-linker-CYFP protein fusion, and a CsrA-linker-NYFP fusion. Direct CsrA-mRNA binding results in complementation of the YFP protein and generates a fluorescence output. c) Ratio of median YFP with the target RNA-MS2BD-terminator fusion relative to *fecA*-MS2BD construct, which was used as a negative control since this target is confirmed to not interact with CsrA. The TriFC components were expressed in *E. coli* Δ*csrB* K-12 MG1655. Means represent five biological replicates and error bars indicate propagated standard deviation. Asterisks indicate a p-value < 0.05 of a heteroscedastic T-test between the fluorescence ratio of the mRNA tested and that of the *fecA* negative control. Figure created with Biorender.com.

From these newly confirmed direct targets, we identified the *uxuB* transcript as most interesting for further characterization as it was the only target of the 9 that has previously shown CsrA-dependent activation *in vivo*^29^. Additionally, the *uxuB* transcript is particularly interesting given its unique number, location, and sequence of predicted CsrA binding sites in this mRNA (**Figure 2a**). Although it is possible that CsrA also heterogeneously regulates the other targets (*yqjE*, r*odZ*, *cmk*, *dps*, *ybal*, *acnA*), we did not investigate this possibility.

**Figure 2.**
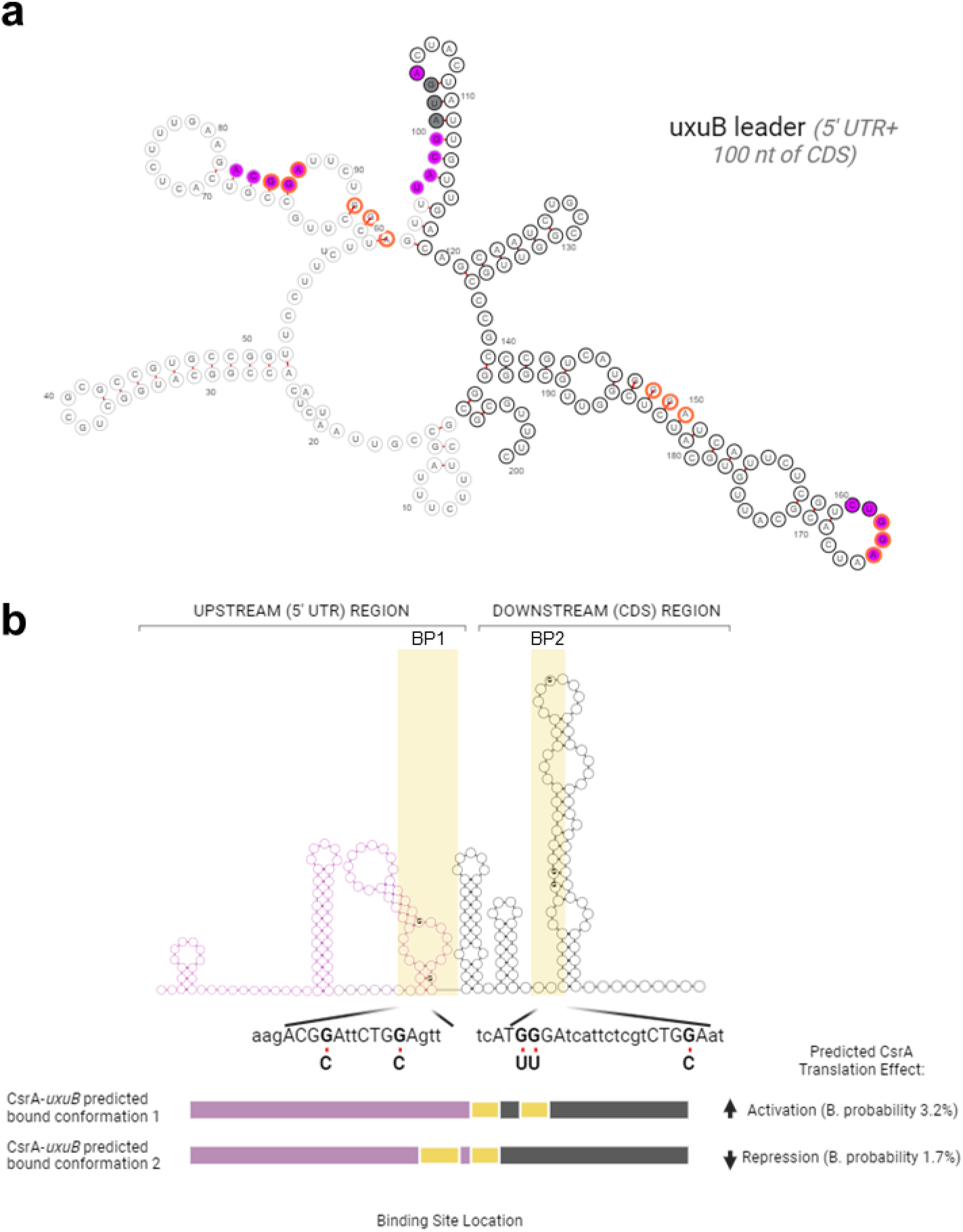
The *uxuB* transcript contains two binding pockets predicted to mediate opposed regulatory outcomes. a) *uxuB* 5’UTR + 100 nt of CDS. There are three bioinformatically predicted binding sites (purple) in the *uxuB* leader sequence. The sequence contains four “GGA” sites (highlighted in orange), two of which were not predicted by the model to be CsrA binding sites. The start codon of the *uxuB* transcript is indicated in gray and nucleotides corresponding to the coding sequence are bolded. b) Linear representation of the *uxuB* transcript. The 5’ UTR of *uxuB* is colored purple and the coding sequence is shown in black. Below is a schematic showing the top two CsrA-*uxuB* bound conformations identified^38^ which result in opposed regulatory outcomes (the values shown in the bottom right side of the figure indicate the Boltzmann probability for these conformations). In these predictions, two binding pockets (BP1 & BP2) contain two high-affinity GGA motifs each and binding at these pockets potentially leads to different regulatory effects in translation of *uxuB* (*i.e*. BP1 is predicted to be used for repression and BP2 for repression).

The CsrA protein binds RNAs as a homodimer, canonically at two stem loop regions containing a high affinity “GGA” motif -with an optimal spacing of 18 nts between binding sites^42^. For *uxuB*, there are four predicted high-affinity GGA motifs that fall within the 5’ UTR through first 100 nts of coding sequence; we define this sequence range as the *uxuB* leader sequence. Importantly, these predicted CsrA-binding GGA motifs are optimally spaced between one another, based on previous *in vitro* binding assays^42^. Additionally, these sites are located at both the 5’ UTR and the coding sequence, which could potentially allow for multiple modes of regulation. Within the *uxuB* leader sequence there is a binding pocket in the 5’ UTR, referred to as Binding Pocket 1 (BP1), which encompasses two GGA motifs at the 5’ UTR of the *uxuB* transcript. Binding within BP1 is computationally predicted to result in translational repression by CsrA^38^ (**Figure 2b**). A second binding pocket, Binding Pocket 2 (BP2), is located in the coding sequence and contains two additional GGA motifs. Binding in this region is predicted to result in translational activation^38^ (**Figure 2b**).

### CsrA interacts independently and simultaneously with two binding interfaces of the *uxuB* transcript

As mentioned above, the *uxuB* leader sequence contains four GGA motifs, which represent potential sites for direct CsrA binding. To investigate how these sites individually contribute to CsrA binding, we simultaneously mutated potential GGA binding sites in two discrete binding interfaces: the 5’ UTR region, which contains BP1 and the coding (CDS) region, which contains BP2 (**Figure 3a**). We then evaluated changes in binding affinities for each mutant via EMSAs (**Figure 3b, c**). When possible, we made a GGA to GCA mutation since this base substitution has been shown to be sufficient to eliminate CsrA binding^43^. We made other conservative mutations when needed, to preserve mRNA secondary structure and the overall base-pairing probability calculated by RNAfold^44^ from the Vienna RNA webserver. For potential binding sites located in the *uxuB* coding region, we also ensured that the mutations introduced were synonymous and did not change the codon identity of individual triplets. The resulting sequences for each mutant version tested can be found in **(Supplementary Table S2)**.

**Figure 3.**
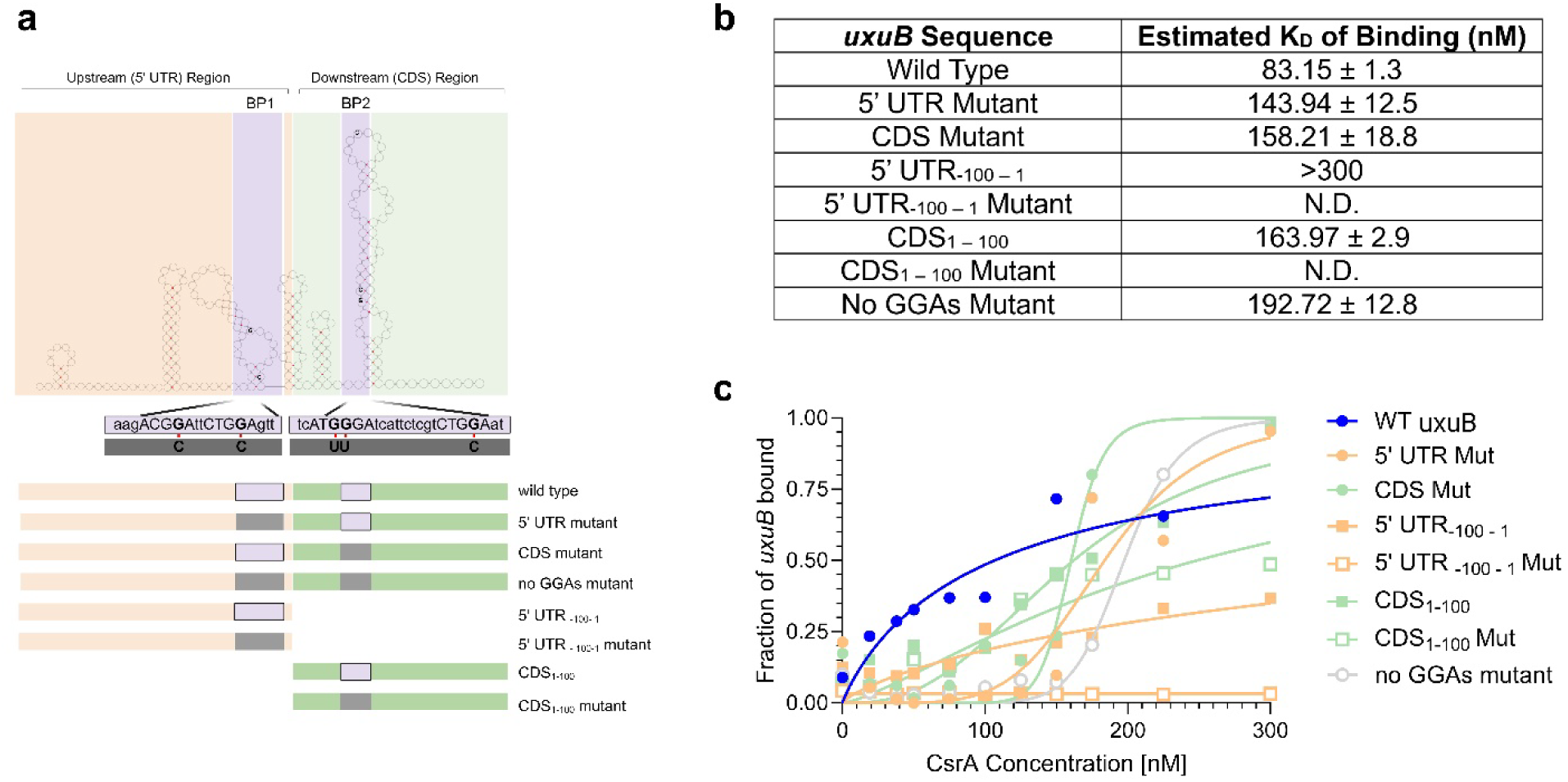
Two binding interfaces in the *uxuB* transcript mediate CsrA binding. a) Linear map of the *uxuB* leader sequence (5’ UTR + first 100 nt of coding sequence). We divided the *uxuB* transcript in two distinct binding interfaces denoted as 5’ UTR region (orange) and CDS region (green) each interface containing potential CsrA-binding sites (purple) depicted as binding pocket 1 (BP1) and binding pocket 2 (BP2). The mutations introduced to eliminate potential CsrA binding at each binding site are outlined and colored grep. b) Table summarizing the equilibrium constant estimates for each *uxuB* variant as determined via electrophoretic mobility shift assays. c) Individual data points and fitted binding curves.

We experimentally determined CsrA to bind *uxuB* with an apparent KD ∼ 83.15 ± 1.3 nM. This value is within the range of previously reported KD values for CsrA binding to other targets in *E. coli* which vary between 4-95 nM for mRNA targets^45,46^. Mutations to both the 5’UTR and CDS regions demonstrated reduced affinity between CsrA and the mutant *uxuB* transcripts, as indicated by the increase in KD values (K1/2 ∼ 143.94 ± 12.5 nM and K1/2 ∼ 158.21 ± 18.8 nM, respectively). CsrA binding affinity for *uxuB* was further reduced when all GGAs were removed from the sequence (K1/2 ∼ 192.72 ± 12.8 nM) **(Figure 3b).** Importantly, upon mutation of one of these interfaces, binding curves adopted a sigmoidal shape, indicating cooperative CsrA binding. Overall, these results suggest that both regions are important and contribute to CsrA-*uxuB* binding.

To investigate whether CsrA could interact independently with each of these binding interfaces, we generated truncated sequences for both the 5’ UTR and CDS regions of the *uxuB* leader sequence (**Figure 3a**). Briefly, these truncations isolated either the 5’ UTR or CDS. Additionally, for each individual region, we made mutations to remove the GGA sites. In these experiments, CsrA bound the CDS truncation with stronger affinity (KD ∼ 163.97 ± 2.9 nM) relative to the 5’ UTR truncation which was bound by CsrA very weakly (KD >300 nM) (**Figure 3b**). Mutations to the GGA sites on each truncated region virtually eliminated binding in the case of the 5’ UTR truncation mutant (**Figure 3c**). This suggested to us a potential preference of CsrA to the binding sites located in the CDS region, which agrees with our previous biophysical model results. Furthermore, the curvature and the estimated Hill coefficient of approximately 2 for the CDS truncation binding curve suggest cooperative binding. Interestingly, a similar behavior was recently reported for truncations of the CsrA target *yqjD* suggesting similar mode of binding^46^. Gel images for each mutant and/or truncated sequence can be found in (**Supplementary Figure S1**). Together, these results indicate that CsrA binds to *uxuB* with similar affinity to previously characterized targets. Additionally, these results suggest that CsrA binds to *uxuB* using both the 5’ UTR and coding regions (CDS) with a stronger preference for the interface located within the CDS of *uxuB*.

### CsrA activates and deactivates *uxuB* expression in a concentration-dependent manner

As mentioned previously, CsrA is predicted to have a heterogenous regulatory effect on uxuB translation. After identifying the regions within the *uxuB* transcript that mediate CsrA binding, we next assessed the regulatory outcome of these interactions. To achieve this, we employed *in vitro* coupled transcription-translation (IVTT) PURExpress system. We chose the IVTT system as it allowed us to evaluate regulatory effects mediated by CsrA directly without additional confounding cellular factors which might contribute to indirect regulation. Our IVTT assays utilized *uxuB- gfpmut3* fusion sequences of the wildtype as well as 5’ UTR and CDS mutant *uxuB* leader sequences cloned into a T7-driven expression plasmids (**Figure 4a)**. Each of the *uxuB* sequences included -100 through the +100 nucleotides, with specific point mutations made in the mutant constructs. Additionally, the full sequence for this construct is included in **Supplementary Table S2.** Importantly, the IVTT reactions were prepared in the absence of CsrA as well as with increasing concentrations of CsrA as previously described^26^. For the wildtype *uxuB*-*gfpmut3* sequence, we observed a 2.1-fold increase in GFPmut3 fluorescence with CsrA at a concentration of 125 nM. At 250 nM of CsrA, we observed reduced signal activation in our IVTT assays, and observed no significant signal activation in all reactions with 375 nM of CsrA or higher **(Figure 4b, left panel)**.

**Figure 4.**
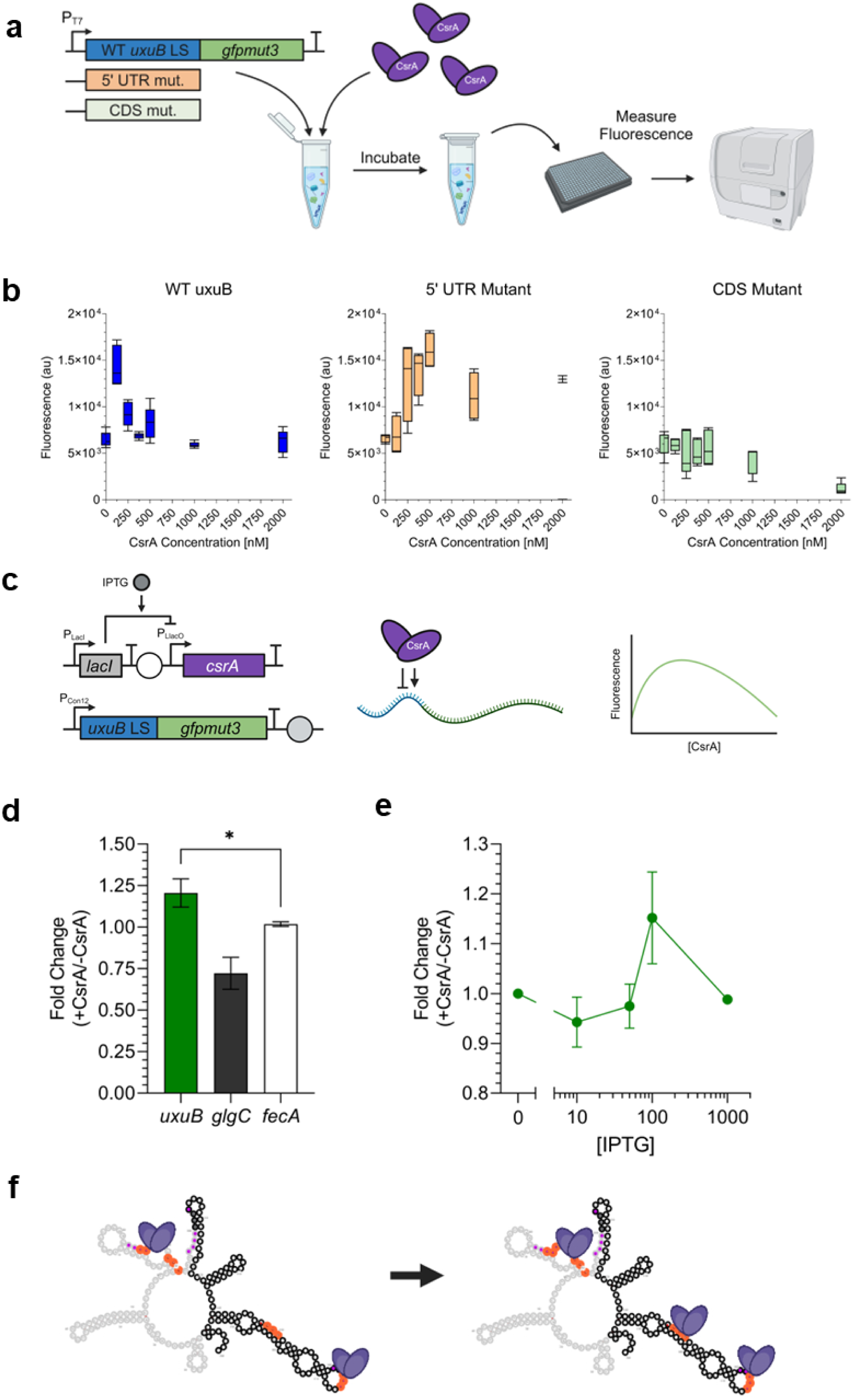
Demonstrating *in vitro* and *in vivo* CsrA dose-dependent regulation of *uxuB* expression. a) Components and workflow of the *In Vitro* Transcription-Translation (IVTT) assays performed to evaluate CsrA-mediated regulation of *uxuB-gfpmut3* fusions. Individual reactions were incubated with fixed concentrations of CsrA (0-2000 nM), and fluorescence was monitored for 3 hours. b) Endpoint GFPmut3 fluorescence for reactions containing wild type (left), CDS mutant (middle), and 5’ UTR mutant (right) *uxuB* leader sequences incubated at increasing concentrations of CsrA. Box plot represents the distribution of the data for technical quintuplets. Significance was evaluated by comparing the fluorescence values of the reactions with CsrA (125-2000 nM) to those of the reaction without CsrA (0 nM) for each *uxuB* leader sequence tested using a heteroscedastic unpaired t-test. Asterisks denote p-values ≤0.05 from this analysis. c) Schematic for *in vivo* reporter assay. The *uxuB* leader sequences fused to *gfpmut3* are constitutively expressed from one plasmid, while CsrA is titrated via IPTG induction from a second plasmid. *In vivo* regulation of the *uxuB-gfp* fusion reporter by CsrA at different concentrations generate fluorescent signals. d) Fold change in GFPmut3 fluorescence for the WT *uxuB* leader sequence, as well as *glgC* and *fecA* leader sequences and positive and negative controls. Fold change is calculated as the fluorescence of each *leader-gfpmut3* 3 hours post-induction induction of CsrA divided by the cultures without CsrA induction, as previously established^29^. e) Fold change in fluorescence of the WT *uxuB-gfp* reporter at increasing concentrations of CsrA. f) Proposed model of dose-dependent regulation of *uxuB* by CsrA. Figure created with Biorender.com.

We also tested a CDS mutant, in which the downstream GGA motifs were mutated such that CsrA can only interact with the remaining binding sites in the 5’ UTR and observed that titrating CsrA could no longer activate GFPmut3 signal (**Figure 4b, middle panel**). These results further support that CsrA binding in the CDS region of *uxuB* mediates target activation. Lastly, we tested a 5’ UTR mutant where the GGA motifs in the 5’ UTR were mutated, leaving only those in the CDS region. For the 5’ UTR mutant, we observed significant GFPmut3 signal activation at 250 nM of CsrA and greater with activation increasing as CsrA concentrations increased (**Figure 4b, right panel**).

We next sought to evaluate CsrA regulation of *uxuB* in an *in vivo* context with a plasmid-based reporter assay previously used by our group and others to evaluate CsrA regulation of target mRNA transcripts^29,47^. Briefly, we fused the *uxuB* leader sequence in-frame to *gfpmut3* and constitutively expressed this transcript from a plasmid, while CsrA is expressed from a second plasmid via IPTG induction (**Figure 4c**). Comparing GFPmut3 fluorescence between induced and uninduced samples are used to determine the regulatory effect upon CsrA binding (**Supplementary Figure 3a**). We performed this experiment in an *E. coli* MG1655 K-12 Δ*csrABCD*Δ*pgaABCD*Δ*glgCAP*, referred to as the Δ*csr* strain henceforth, such that our plasmid-based CsrA is the only component from the Csr Network interacting with the *uxuB* transcript. The *glgCAP* and *pgaABCD* operons are deleted to ensure cellular fitness upon deletion of *csrA*^47^.

In our *in vivo* assay, we observed that cultures induced with 100 µM of IPTG resulted in significant signal activation from the *uxuB-gfpmut3* construct relative to our negative control (**Figure 4d**). We use the leader sequences of *fecA* as a negative control and the *glgC* leader sequence as a positive control as previously done^29^. The leader sequence of *glgC* is a well-characterized target repressed by CsrA^48^ and was thus used as positive control. Additionally, our results are in accordance with previous reporter assays that demonstrated CsrA-mediated activation of *uxuB*^29^. Interestingly, at lower and higher concentrations of CsrA induction, we observed no signal activation of the *uxuB* reporter (**Figure 4e**). When we tested the *uxuB* 5’ UTR mutant (CsrA can only bind the CDS) in our plasmid-based assay we observed target activation suggesting that CsrA binding in the coding region of *uxuB* leads to activation. The opposite effect was seen when testing a *uxuB* CDS mutant (CsrA can only bind the 5’ UTR), indicating that CsrA to the 5’ UTR of *uxuB* leads to target repression (**Supplementary Figure S3**).

These collective results continue to support our proposed model of dose-dependent CsrA regulation of *uxuB.* We interpret these results as CsrA interacting first with a preferred CDS region of *uxuB* at low intracellular CsrA concentrations. At high CsrA concentrations, additional interactions could occur in the 5’ UTR region of *uxuB*, resulting in deactivation of *uxuB* expression. Our working model of this concentration-dependent regulation of *uxuB* by CsrA is illustrated in **Figure 4f**. Physiologically, these two modes of regulation are possible since free intracellular CsrA is sequestered by the CsrB and CsrC sRNAs under low glucose availability^49^ and activating the *uxuAB* operon under these specific environmental conditions would allow the cell to metabolize secondary carbon sources such as D-glucuronate or D-fructuronate. Moreover, CsrA concentrations *in vivo* reportedly vary approximately 2-3-fold during different growth phases. Additionally, its binding activity slightly increases between lag and exponential phase, while greatly increases between exponential and stationary phase^50^. These results also support our hypothesis of dose dependent heterogenous regulation, as a slight increase in CsrA binding activity would activate utilization of the *uxuAB* operon to metabolize secondary carbon sources during exponential phase, then deactivate those pathways upon entering stationary phase due via increased binding activity.

Notably, when evaluating the impact of CsrA regulation of *uxuB* on secondary metabolism, we did not see significant changes in hexuronic acid utilization, concretely demonstrated by the lack of any visible growth changes between WT MG1655 and a genomic *uxuB* mutant lacking all GGA (*no_GGAs*) when grown on only glucose, only glucuronate, or glucose plus glucornate (**Supplementary Figure S4**). This observation suggests that CsrA regulation of this pathway may be more complex than what we are able to replicate under standard laboratory conditions.

### Accessibility probing reveals dose-dependent structural rearrangement of *uxuB*

The concentration-dependent effects on *uxuB* expression (**Figure 4**) led us to hypothesize that CsrA binding may cause structural rearrangements to the *uxuB* leader sequence that initially promotes cooperative binding of additional CsrA dimers. A similar model has recently been proposed for the binding of CsrA to the *yqjD* leader sequence^46^.

To assess dose-dependent changes in regions on the *uxuB* transcript assumed to be available for CsrA binding, we adapted a cell-free antisense probing approach (abbreviated CF-iRS^3^)^51^. This method relies upon the conformational change of a toehold switch that is induced by RNA-RNA base pairing between an antisense probe and an accessible region of the RNA target. In this scheme, RNA bases that are unpaired are more accessible for interacting with the antisense probe and yield a higher fluorescent signal than those that are blocked by either the targets’ secondary structure or by protein binding. (**Figure 5a)** We used this approach to probe several predicted CsrA binding sites on the *uxuB* leader sequence as well as other regions of the transcript as a control; probed regions are shown in **Figure 5b** and span model-predicted binding sites (pink) and other potential regions given the presence of high-affinity GGA motifs (orange). Importantly, our probing analysis revealed several key regions change in accessibility with increasing concentrations of CsrA. For instance, Probe 1 (nts -25 to -12), which spans the first predicted binding region in the 5’ UTR of *uxuB* is located upstream of the ribosome binding site and demonstrated significant increase (p < 0.05) in accessibility in the presence of 125 nM CsrA relative to accessibility at 375 nM CsrA (**Figure 5b**). The site Probe 1 occupies is upstream of the ribosome binding site (within 5’ UTR region) and is inaccessible in the absence of any CsrA (*uxuB* only), which suggests that binding influences a structural rearrangement in this region. Probe 2 (nts -14 to +3) spans the remainder of the *uxuB* 5’ UTR covering the ribosome binding site and start codon. Probing this area showed a significant (p < 0.05) reduction in accessibility in the presence of 375 nM of CsrA but remained unchanged at the lower 125nM CsrA concentration relative to baseline accessibility of *uxuB* with no CsrA present (**Figure 5c)**. This suggests that CsrA binds to the ribosome binding site and start codon region only after higher affinity sites are occupied. Probes 3, 4, and 5, are all within the CDS region of the *uxuB* leader sequence (**Figure 5b)**. Probe 3 (nts +36 to +51) spans a region not predicted to be bound by CsrA, but still contains a high affinity GGA motif. Interestingly, accessibility was significantly elevated in this region in the presence of both concentrations of CsrA (p < 0.05) (**Figure 5c**). This suggests that CsrA is not directly interacting with this site, but that binding in other regions may be causing the structure surrounding this region to open. We did not observe any significant changes in fluorescence for Probes 4 (nts +52 to +66) and 5 (nts +73 +86) when incubated in the presence or absence of CsrA (**Figure 5c**). The lack of detectable change at Probe 4 may have been caused by low accessibility of the region to begin with, as evidenced by the low fluorescent signal measured in the absence of CsrA (*uxuB* only control). The lack of changes in accessibility, as measured by lack of fluorescence changes, was expected for Probe 5 as we selected this region to act as a negative control for CsrA binding.

**Figure 5.**
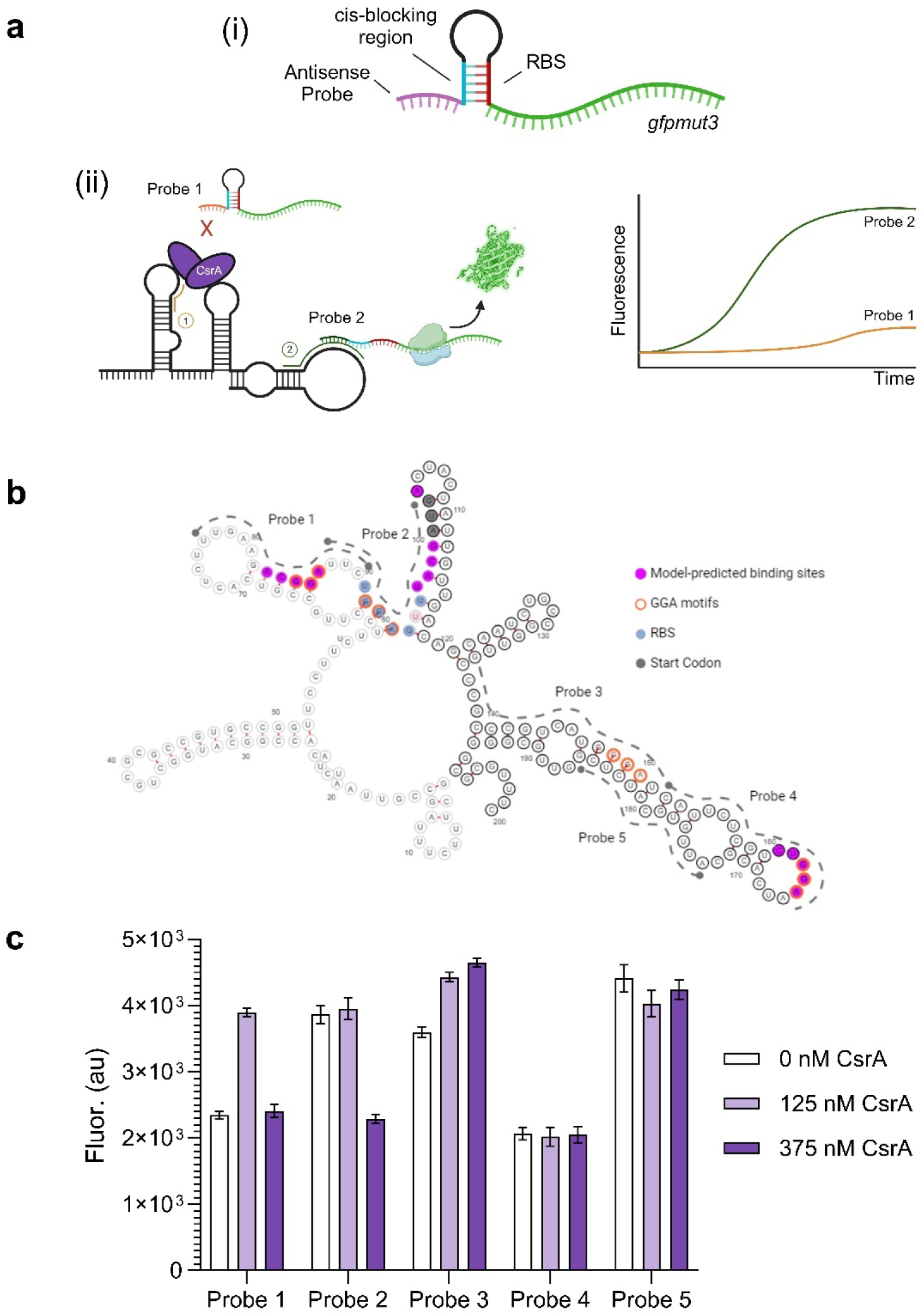
***In vitro* antisense probing of *uxuB* RNA accessibility reveals structural changes in the presence of low and high concentration CsrA.** a) i. Diagram of the CR-iRS^3^ probing structure. We use a short anti-sense probe to bind to regions of interest along an RNA. ii. When a probe cannot access its target region (Probe 1), the RBS remains sequestered, and minimal signal is observed. When a probe successfully binds its target (Probe 2), the hairpin unwinds and allows translation to occur, generating a signal. b) Predicted structure of *uxuB* RNA used for this assay. Regions probed using CF-iRS3 assay are labeled in dashed lines with genomic coordinates. Areas with predicted binding sites (pink) and GGA motifs (orange) are highlighted, along with key functional regions of the leader sequence. Samples containing *uxuB* leader RNA were incubated with 0, 125, and 375 nM CsrA protein and probed using CF-iRS^3^ antisense probes designed for specific regions 1-5. c) Endpoint GFPmut3 signal of each probe region in presence of 0, 125, and 375 nM CsrA.

Collectively, our interpretation of these results is that CsrA binding to higher affinity sites within the CDS region of the *uxuB* leader sequence allows for multimerization of CsrA proteins on the *uxuB* transcript, preventing translation. These results, along with intermediate bands observed in mobility shift assays (**Supplementary Figure S1**), appear to be consistent with our proposed mechanism. Direct binding between CsrA and the 5’ UTR has been shown to induce structural rearrangements that allows for increased ribosomal binding, leading to increased translation; this has been observed in the activating effects of CsrA binding to the *ymdA* 5’ UTR^52^ and RsmA binding to the *phzA2* 5’ UTR^53^. In addition, hierarchical binding that allows for multimerization is observed within the CsrA ortholog RsmE and the sRNA sponge RsmZ in *P. fluorescens*^54^. In these cases, binding of initial Csr/Rsm proteins to high affinity sites results in structural rearrangement within the target RNA that allows for more Csr/Rsm proteins to bind to a single transcript.

This hierarchical binding strategy may be leveraged by CsrA within *uxuB* to result in a dose-dependent switch in regulatory outcomes. Specifically, a hierarchical strategy in which CsrA first binds to high affinity sites that results in a structural rearrangement surrounding the RBS which results in activation of expression. The RBS, containing a high affinity A(N)GGA motif, is then available to be bound by CsrA which switches the system towards a repressive state. This reflects a novel strategy of regulation within the 5’UTR of an mRNA which contain core conserved regions surrounding the RBS that can shift regulatory outcomes. Overall, the shift in accessible regions identified by our probing approach suggests that such the phenomenon of dose-dependent multimerization may be occurring.

### Leveraging the dose-dependent response to construct a minimal BANDPASS gate

Based on the unique dose-dependent response between *uxuB* and CsrA, we sought to generate complex post-transcriptional synthetic regulators. The dose-dependent response cycle of no signal to activation back to no signal closely follows that of a BANDPASS filter architecture. The BANDPASS filter is derived from electrical engineering principles, which a circuit achieves signal amplification only within a specific range of input frequencies^55^. In a biological engineering context, this can be translated to a genetic circuit that only activates gene expression of its target within a specific range of input signal (i.e., small molecule inducer). Biologically-derived BANDPASS filters have previously been engineered using transcriptionally regulated components^56–59^.

In our own recent work^27^, we developed synthetic circuit architectures to leverage native CsrA-5’ UTR interactions, such that 5’ UTRs can be “drag-and-dropped” into a circuit and recapitulate the native CsrA-RNA regulatory outcome. Here, we adapt this system to design a new Csr-regulated BANDPASS filter by fusing the -100 through the +100 nucleotides of the *uxuB* mRNA transcript directly upstream of the *gfpmut3* CDS (**Figure 6a, top**). The native start codon within *uxuB* was mutated from ATG to AAA to eliminate any ribosome competition for the *gfpmut3* start codon. Using the ATG to AAA mutant, the Csr-regulated BANDPASS filter is established as follows: (i) in the absence of IPTG, the filter begins in the “OFF” state as intracellular CsrA interacts with the *uxuB* sequence to prevent translation of GFPmut3, (ii) the CsrB sRNA is expressed from the plasmid as IPTG is titrated into the system and sponges away a portion of intracellular CsrA to reduce intracellular CsrA concentrations, and eventually sufficient CsrA is sequestered by CsrB such that translation activation occurs establishing the “ON” state, (iii) as CsrB sRNA transcription saturates, there is insufficient intracellular CsrA required to activate translation, returning the system to the “OFF” state (**Figure 6a, bottom**).

**Figure 6.**
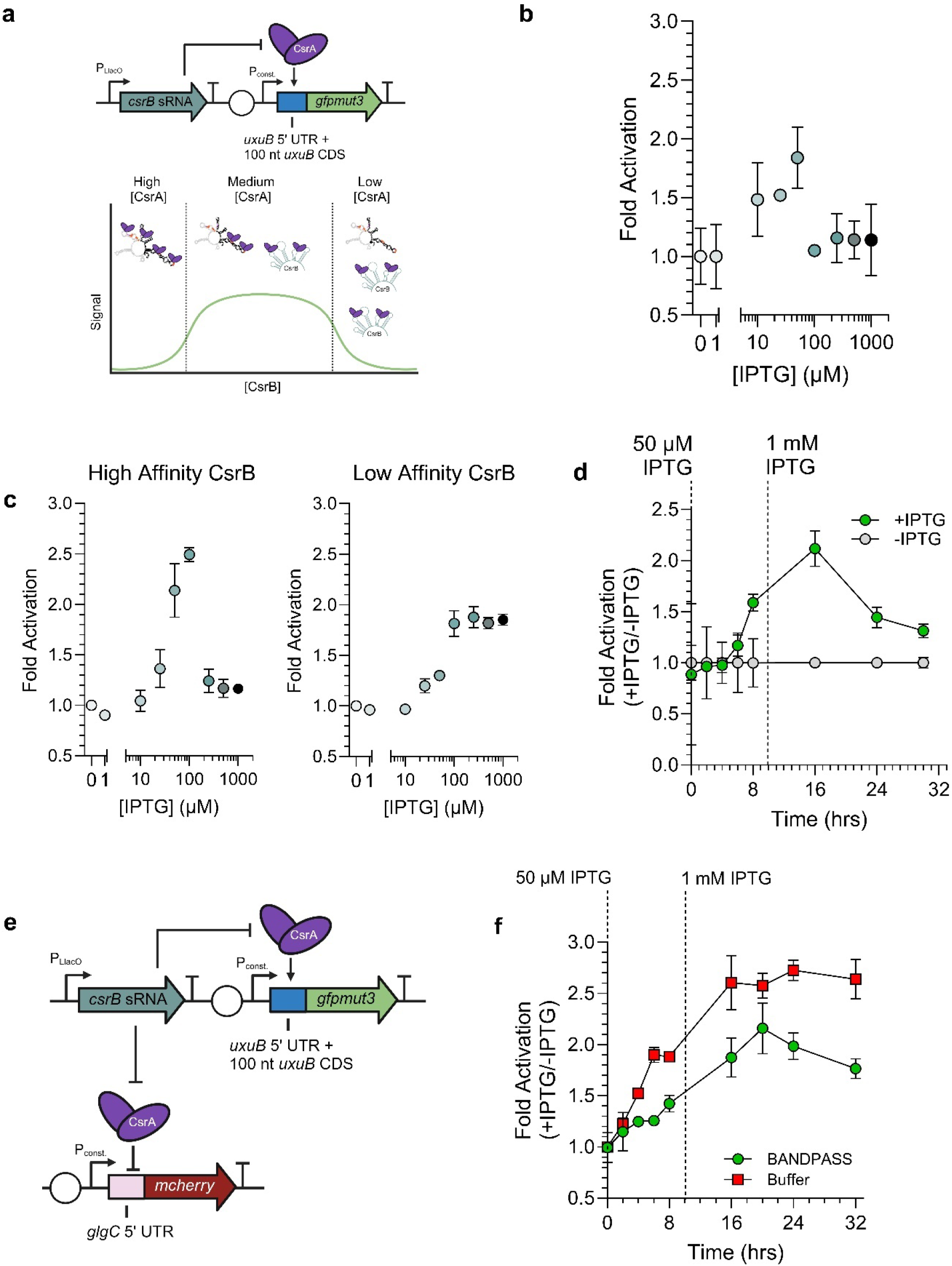
Repurposing the *uxuB*-CsrA Interaction to create a post-transcriptional BANDPASS Filter. a) Genetic circuit design of the Csr-regulated BANDPASS filter (top), and a schematic of the expected outcome of the post-transcriptional BANDPASS filter. b) BANDPASS filter response measured by fold activation of GFPmut3 fluorescence relative to the uninduced control at increasing IPTG concentrations. This BANDPASS filter used the wild type sequence of the CsrB sRNA. c) BANDPASS Filter response for the H11 CsrB (High Affinity) and L2 CsrB (Low Affinity) sRNAs. d) Time course response of the BANDPASS Filter. At seeding the filter was induced with 50 µM IPTG to activate the filter, then fully saturated with IPTG at 1000 µM ten hours post-seeding to deactivate the filter. e) Genetic circuit design to integrate the Csr- regulated BANDPASS filter with the Csr-regulated BUFFER Gate (cBUFFER). f) Time course response of the BANDPASS filter regulating GFPmut3 and cBUFFER regulated mCherry logic measured by fold activation of each fluorescent signal.

To evaluate the dynamics of this BANDPASS circuit design, we titrated CsrB expression and measured GFP fluorescence 12 hours post-induction. For IPTG concentrations less than 10 µM IPTG and greater than 100 µM, we observed residual signal activation. For concentrations between 10 µM and 100 µM, however, we observed signal activation with maximum activation at 50 µM of IPTG (**Figure 6b**). These results demonstrate that the native dose-dependent response between the *uxuB* transcript and CsrA can be leveraged to create a proof-of-concept BANDPASS Filter. In comparison to a previously established BANDPASS Filter^59^, our system demonstrates similar signal activation (**Supplementary Figure S5**), as measured by a similar 2-fold increase in signal expression in each system.

With this circuit established, we sought to tune the response of the BANDPASS Gate by replacing the wild type CsrB sRNA with mutant sequences that were previously demonstrated to alter the affinity towards CsrA^27,37^. Specifically, we tested two CsrB sRNA mutants termed H11 and L2^37^, which previously showed a 3.5-fold and 0.75-fold change in affinity towards CsrA compared to that of the wild type CsrB sRNA. Using these mutant CsrB sRNAs we observed that integrating the high affinity H11 CsrB increased the maximum signal response to 2.5-fold and shifted the activation to occur between 25 and 100 µM IPTG. Integrating the low affinity L2 CsrB shifted activation to only occur above 100 µM of IPTG (**Figure 6c**), characteristic typical of a Buffer Gate. Next, we attempted to leverage the BANDPASS Filter to activate then eliminate GFP expression by increasing the final concentration of IPTG in the system over time. To understand dynamics of GFP expression from the BANDPASS Filter, we tracked signal activation of cultures containing the BANDPASS Filter induced at 50 µM IPTG over a time interval of 30 hours and observed significant signal activation five hours post-induction (**Supplementary Figure S6**) before reaching maximum activation at 24 hours post-induction. With these dynamics in mind, we designed an assay in which we induced WT CsrB expression in the BANDPASS filter immediately after seeding with 50 µM IPTG tracked fluorescence over time. As expected, we observed signal activation relative to the uninduced cultures six hours post-induction. Four hours later, we added more IPTG into the system bringing the final concentration of the culture to 1000 µM IPTG. Interestingly, we observed GFP fluorescence peak at two-fold activation 16 hours after seeding before decreasing to 1.3-fold activation relative to the uninduced samples (**Figure 6d**). We suspect the samples containing the induced BANDPASS Gates do not fully return to baseline due to the stability of the GFP protein.

Lastly, we sought to utilize the BANDPASS filter in parallel with another Csr-regulated logic gate to achieve multiple regulatory outcomes. Specifically, we wanted to evaluate if we could activate two circuits in one environmental condition, then in a second condition inactivate the BANDPASS while the other circuit remains active. To test this, we employed a Csr-regulated BUFFER Gate (cBUFFER) controlling mCherry expression in tandem with our BANDPASS filter. In these experiments, we transformed our BANDPASS Filter regulating GFP concurrently with a Csr Buffer Gate regulating mCherry (**Figure. 6e**) previously developed in our lab^27^. We induced CsrB expression upon seeding with 50 µM IPTG to activate both Gates and observed fluorescence of GFP and mCherry eight hours post-induction. After 10 hours, we then added more IPTG such that the final concentration reached 1000 µM. We observed the Buffer Gate increase over the course of the next six hours and saturate at a 2.5-fold increase from an uninduced control, while the BANDPASS Filter peaked at 2.1-fold increase in signal activation relative to an uninduced control after 20 hours from initial seeding, before decreasing to a 1.7-fold increase after 32 hours (**Figure. 6e**). Overall, results from these experiments provided preliminary that the BANDPASS filter can be used in tandem with other gates. To our knowledge, this is the first demonstration of an RNA-Protein-derived BANDPASS gate demonstrated in a bacterial system as well as the first instance of integrating a BANDPASS filter with multiple genetic circuits simultaneously.

Overall, in this work we demonstrate that discovery and characterization of novel RNA-Protein interactions is important for enhancing synthetic tools for controlling gene expression in dynamic ways. Although *uxuB* had previously be identified as a putative CsrA-activated target^29,38^, its *in vivo* regulation and interactions with CsrA were yet to be fully elucidated. In this work, we elucidated interactions between the *uxuB* transcript and CsrA and characterized a potential mechanism by which the CsrA protein exerts dose-dependent regulation of this mRNA transcript, resulting in multiple modes of regulation. Specifically, we show that translation of the *uxuB* transcript is activated both *in vitro* and *in vivo* by CsrA only within a certain concentration range of intracellular CsrA. With this knowledge, we demonstrate that the native interaction between CsrA and *uxuB* could be repurposed to engineer a tunable post-transcriptional BANDPASS filter that reduces the number of synthetic parts required, compared to previous biological BANDPASS filters, and can be integrated with other post-transcriptional logic gates to achieve complex genetic regulation.

One advantage of our post-transcriptional BANDPASS filter design is that it only requires three synthetic components; the LacI regulator, the CsrB sRNA and the *uxuB* sequence fused to the gene of interest. In contrast, other previously developed systems require multiple nested regulatory components, as well as genome engineering to achieve a BANDPASS response.^59^ Additionally, a total of 5 synthetic parts are required, while still achieving similar fold activation responses (**Supplementary Figure S5).** As our system utilizes less parts, we were able to achieve circuit compression of approximately 1.66-fold. Other BANDPASS filters have managed to achieve greater signal amplification, however their work relied on a 6-component system integrated into the genome, as well as induction by H2O2 . While the signal amplification achieved is impressive, the requirement of genome editing reduces its portability and its reliance on H2O2 as an inducer may further trigger cellular burden and/or oxidative stress^60^.

Another advantage of this gate is the possibility of modifying it to rely on other common small molecule inducer schemes (aTc, vanillic acid, etc.) due to its design where only CsrB must be synthetically regulated. Reducing the synthetic components is advantageous in genetic circuits as it can both reduce cellular burden, and the potential for system failure, which is commonly observed as system complexity increases^13,19,57^. This circuit is also fully self-encapsulated and relies on native CsrA expression to achieve regulation. In this sense, compression of our BANDPASS Filter eliminates the need for nested regulatory components and reduces cellular burden by not relying on high concentrations of antibiotics to attenuate signal amplification. Moreover, we demonstrate that by varying the CsrA-CsrB affinity within the BANDPASS Gate we can tune the environmental conditions under which signal amplification occurs (**Figure. 6c**). That is, this circuit provides temporal control of genetic regulatory outcome that is directly tied to a single input (i.e. variation of CsrB concentrations via changes in an inducer like IPTG), as titrating the concentration of IPTG in the cultures can activate or repress the target gene outcomes based on the CsrB transcribed from the BANDPASS Filter. The latter represents a much more direct approach to tuning the BANDPASS Filter dynamics. Many previously engineered bacterial BANDPASS filters require multiple input signals or a larger number of regulatory components to achieve similar outcomes^59,61^. It is also worth noting that this synthetic system can also remain in one regulatory state for at least 24 hours (**Supplementary Figure S5**), emphasizing that the regulatory interactions of the BANDPASS filter are maintained across several growth stages of the culture.

An exciting aspect of the resulting BANDPASS filter is that it sets the foundation for potential application in dynamic metabolic control (DMC) applications. DMC is a bioprocessing approach, in which genetic circuit activity is coupled with cell growth^62^. Coupling cell growth or metabolism with the synthetic circuits allows for the microbes to dynamically tune heterologous gene expression in response to environmental signals (stressors, nutrients, etc.). We envision that our BANDPASS filter could be utilized to fine tune metabolic pathways that generate toxic intermediate, such that the BANDPASS filter can maintain a steady-state concentration of our toxic intermediate that supports production of the final metabolite without causing additional stress. Moreover, our efficient design will be able to reduce the burden placed on the cells by the BANDPASS filter itself compared to other designs.

## Materials & Methods

### Plasmids and strains

*E. coli* strains used in this study are listed in Supplementary Table S1. The cloning strategy used to generate plasmids as well as detailed description of each construct used in this study can be found in Supplementary Table S1. Oligos used as templates for T7-driven transcription of RNA fragments and cloning were purchased from Integrated DNA Technologies (IDT) and are listed in Supplementary Table S2. Plasmid sequences were verified by long-read sequencing (Plasmidsaurus). The pUC19-T7link-sfGFP parent plasmid used for cloning and *in vitro* transcription-translation assays was a generous gift from Dr. Svetlana Harbough at AFRL. The no_GGAs mutant, the strain in which the CsrA binding sites were mutated in the *uxuB* leader sequence, was synthesized using CRISPR-Cas9-lambda red gene editing protocols previously established^26,63,64^. In short, a plasmid constitutively expressing WT Cas9, and arabinose-inducible lambda red recombinase proteins was transformed into WT K-12 MG1655 *E. coli*. Single colonies were grown overnight in 5 mL of LB supplemented with 50 µg/mL Carbenicillin using a shaking 37 °C incubator. The overnight was used to see a 50 mL LB + carbenicillin culture 1:100, which was grown for 1 hour in the same 37 °C shaking incubator. The culture was induced with 1% w/v L-arabinose, then grown until the OD600 reached 0.4-0.6. At that point, the culture was concentrated and made into electrocompetent cells. A single 100 µL of electrocompetent cells were transformed with a chloramphenicol-resistant plasmid constitutively expressing a gRNA sequence that targeted the *uxuB* sequence in the genome, as well as a gBlock that contained the *uxuB* leader sequence with the GGA sites mutated. The targeting site was selected using the CRISPR gRNA Design webtool from Atum (atum.bio). Cells were recovered for 3 hours in 1 mL LB supplemented with 1% w/v arabinose, then plated on LB Agar plates supplemented with carbenicillin and chloramphenicol. The genomic *uxuB* region from single colonies were amplified using cPCR, then the fragments were purified using a PCR Clean-up kit (Qiagen), before being submitted for sequencing (Plasmidsaurus) to confirm integration of the mutant *uxuB* leader sequence. Successful mutants were then cured of the plasmids according to the protocols previously established^26,63,64^.

### *In vivo* three-component fluorescence complementation assays (TriFC)

Fluorescence assays were performed as described in^40,41^ with slight modifications. The two-plasmids system (pTriFC: RNA-MS2 binding domain-*rrnB* fusion + CsrA-NYFP & pMS2-CYFP: MS2-linker-CYFP) was transformed into *E. coli* MG1655 K-12 Δ*csrB*. Biological quintuplet colonies were picked and grown to saturation in 5 mL cultures at 37°C overnight. Fresh 5 mL subcultures were seeded (1:100 dilution) and grown at 37°C until mid-exponential (OD600∼0.4- 0.6). Expression of the fusion constructs from pLacO promoters was induced with 1 mM IPTG (final concentration). Cultures were allowed to grow at 37°C for a total of 22 hours. Yellow fluorescence was measured in a LSRFortessa Flow Cytometer (BD Biosciences) and median fluorescence values were computed. Fluorescence medians were compared to that of the *fecA- MS2-rrnB* fusion negative control. Statistical significance was determined using two-tailed heteroscedastic t-tests.

### *In vitro* RNA-protein binding assays

RNA sequences for mRNA leader sequences (annotated 5’ UTR + first 100 nt of coding sequence) were transcribed using the MEGA Script IVT Kit (Thermo Fisher Scientific) following manufacturer instructions. Double-stranded DNA templates were removed by DNase I digestion and the quality of the transcripts was assessed in 8% urea gels stained with Sybr Green II (Thermo Fisher Scientific). 5’-end labeling was performed using T4 Polynucleotide Kinase (NEB) and [gamma-^32^P] ATP (PerkinElmer). RNAs were purified using DTR Gel Filtration Cartridges (EdgeBio) and their concentration was measured via fluorometric quantification (Qubit RNA High Sensitivity Kit).

Electrophoretic mobility shift assays (EMSAs) were performed as described previously^26^. For initial screening of in vitro binding, 10 nM of radiolabeled RNAs were incubated with 0, 3 (300:1 ratio) or 6 (600:1 ratio) µM of purified CsrA. These ratios were selected based on previously reported affinities for CsrA targets presumed to be physiologically relevant according to CsrA CLIP-seq experiments^28^. All binding reactions were carried out with a large excess of yeast total RNA to inhibit non-specific CsrA-RNA association^45^. For subsequent EMSAs to determine KD values and evaluate the contribution of predicted binding sites to complex formation, additional CsrA concentrations were tested to expand the range in which a CsrA-RNA bound complex will initially be observed (0-300 nM). Radiolabeled RNAs were incubated with purified CsrA for 30 min. prior to loading and running on a 10% non-denaturing polyacrylamide gel with 0.5X TBE running buffer (IBI Scientific, 10x composition: 89 mM Tris, 89 mM Boric Acid, 2 mM EDTA) at 170 V for 6-8 hours at 4 °C. Gels were exposed overnight on phosphor-imaging cassettes (bioWORLD) and imaged on a Typhoon FLA 700 (GE Health Life Science) at 1000 V.

### *In vitro* coupled transcription-translation assays

*In vitro* coupled transcription-translation assays were performed using the PURExpress Kit (New England Biolabs). Template plasmids utilized a *uxuB-sfGFP* fusion sequence. All reactions were performed according to the protocols previously utilized to evaluate CsrA interactions with mRNA sequences^26^.

### Detecting accessible regions of *uxuB* RNA using cell-free toehold switch probing

Accessible regions of the *uxuB* transcript were probed using the CF-iRS3 methodology previously described^51^ with modifications to assess the influence of a binding on regions of the transcript. The *uxuB* leader and first 100 bases of coding sequence was transcribed using the same *in vitro* transcription methodology described in “*In vitro* RNA-protein binding assays’’ and purified using the Zymo RNA Clean and Concentrator spin-column extraction kit (Zymo Research). RNA concentration was then quantified using the NanoDrop® ND-1000 UV-Vis Spectrophotometer. Probe plasmids were constructed using golden gate cloning of pre-hybridized primers (Table S2) and the BsmBI digested backbone as described in^41,51^.

The probing assay was then performed using 1.36 ug of purified *uxuB* mRNA and the addition of 0, 125, and 375 nM of purified CsrA protein. No RNA and no protein conditions were included as negative controls. Previously, the maximum concentration of RNA produced by the PUREexpress system was approximated to be 1- 1.5 uM^65^ which we pre-loaded to approximate stoichiometric parity between the target RNA and the maximum expression T7-transcribed probe mRNA. The maximum CsrA protein concentration of 375 nM was used to reflect the minimum concentration needed to observe repressive effects in the coupled transcription-translation assays. For assay assembly, the transcribed *uxuB* transcript was denatured at 95C for 5 minutes and snap cooled on ice to ensure native folding conformation prior to loading. All reactions were performed in duplicate using PUREexpress *in vitro* protein synthesis reagents (New England Biolabs) supplemented with 1 uL of Murine RNase inhibitor (NEB) and 125 ng of the probe plasmid DNA as per the kit protocol. Purified CsrA protein and *uxuB* RNA were incubated for pre-binding in the PURE reaction mixture at 37C for 30 minutes. 125 nM of the probe plasmid was then spiked in prior to measurement of sfGFP signal production.

sfGFP measurements were captured every 10 minutes on the Cytation3 plate reader (BioTek) for 180 minutes at 37C, with excitation and emission spectra at 470 and 510 nm respectively. Final data analysis was performed using the final fluorescence measurements at steady state.

### *In vivo* fluorescent reporter assays

Fluorescent reporter assays were performed as previously described^26,29^ with slight modifications. The two plasmids (pHL1765, pHL600) were transformed into a K-12 MG1655 Δ*csrABCD*Δ*pgaABCD*Δ*glgCAP* strain of *E. coli* referred to as the Δ*csr* strain. Single colonies were grown overnight in 5 mL of Luria Broth (LB) supplemented with Carbenicillin and Kanamycin at 50 µg/mL in a 37 °C shaking incubator (New Brunswick Scientific I26). Fresh 5 mL cultures of LB supplemented with antibiotics were seeded with 50 µL of saturated overnight culture. Cultures were grown until the OD600 reach approximately 0.2-0.3. Cultures were then induced with IPTG (Research Products International) at various concentrations (0-1000 µM). Cultures were grown for an additional 3 hours after induction in the 37 °C shaking incubator. After the three hours, 1 mL of sample was aliquoted into 1.7 mL microtubes (Olympus) and centrifuged for 5 minutes at 5000 rpms. Liquid waste was aspirated and discarded. The remaining pellet was resuspended in 200 µL of 1x Phosphate Buffer Saline, which was prepared according to Cold Springs Harbor Protocol^66^. The resuspensions were pipetted into a Greiner µClear, F-Bottom, non-binding 96- well plate (Item No. 655906). GFPmut3 fluorescence was measured on a CyTek3 Plate Reader using an excitation wavelength of 488 nm and emission wavelength of 513 nm. OD600 was measured using an absorbance wavelength of 600 nm. 200 µL of 1x PBS was used as sample blanks. All samples were done in biological triplicate.

### BANDPASS Filter and Synthetic Post-Transcriptional Circuit Assays

Experiments to evaluate the post-transcriptional BANDPASS filter were adapted from previous work^27^. Plasmids containing the filter were transformed into K-12 MG1655 *E. coli* competent cells. Single colonies were grown overnight in 600 µL of MOPS EZ-Rich Defined Medium (Teknova) supplemented with 50 µ/mL carbenicillin overnight at 37 °C in a 96-well 2 mL deep-well plate (VWR) shaking at 1100 rpms. The plate was covered with the “Breathe Easier” sealing membrane (MilliPore Sigma Item No. Z763624). The next day 488 µL of MOPS EZ-Rich Defined Media supplemented with antibiotics was inoculated with 12 µL of saturated overnight. Samples were grown at 37 °C in a 96-well 2 mL deep-well plate shaking at 1100 rpms until the OD600 reached 0.2-0.3. The samples were induced with various concentrations of IPTG and then grown for another 12 hours. 50 µL of each sample was then diluted into 150 µL of 1x PBS and transferred into a Greiner µClear, F-Bottom, non-binding 96-well plate. Fluorescent measurements of GFPmut3 were measured on a CyTek3 Plate Reader using an excitation wavelength of 488 nm and an excitation wavelength of 513 nm. OD600 was measured via absorbance with a wavelength of 600 nm Blanks were prepared by adding 50 µL of MOPS EZ-Rich Media to 150 µL 1x PBS. All samples were done in biological triplicate.

For time course experiments, overnight samples were prepared in the same fashion as above. To measure the time course, a sample for each timepoint was prepared as stated above. Additionally, IPTG was added to half of the samples to a final concentration of 50 µM upon seeding. Fluorescence of GFPmut3 and OD600 was measured as before. 10 hours after induction, more IPTG was added to the samples such that the final concentration was 1 mM. Fluorescence and OD600 was monitored for another 20 hours after final induction. For samples with both the BANDPASS filter and the Csr-regulated BUFFER GATE (cBUFFER), the same experimental setup was used, however 50 µg/mL kanamycin was also supplemented to select for the cBUFFER-containing plasmid as well. mCherry fluorescence was measured using the excitation wavelength of 585 nm and the emission wavelength of 613 nm.

To compare the Csr-regulated BANDPASS filter to other biological BANDPASS filters, we synthesized the transcriptional BANDPASS filter previously created^59^. To measure the activity of the transcriptional BANDPASS filter, the protocol established in that work was followed exactly as previously reported^59^.

## Author Contributions

Designed research: A.M.R.N., T.R.S., A.J.L., R.B., and L.M.C.; performed experiments: A.M.R.N., T.R.S., A.J.L., R.B., J.S.R., J.L.A.; analyzed data: A.M.R.N., T.R.S., A.J.L., R.B. and L.M.C.; wrote paper: A.M.R.N., T.R.S., A.J.L., R.B., and L.M.C.

## Notes

The authors declare no competing financial interests.

## Acknowledgements

The authors would like to thank Dr. Tony Romeo, Dr. Han Lim, Dr. Brian Pfleger and Dr. Svetlana Harbaugh for their gifts of several *E. coli* strains and plasmids used in this work. We additionally thank Dr. Abigail Leistra and Dr. Mia K. Mihailovic for insightful discussions and feedback for experimental design. Lastly, we thank Faith Guice and Sung H. Jung for their assistance in preliminary *in vivo* reporter assays. This work was supported by the National Institutes of Health [grant number R01GM135495 to L.M.C., and R.B.]; National Science Foundation [grant numbers MCB-1932780 to L.M.C., DGE-1610403 to T.R.S, and A.J.L..]; the Welch Foundation (grant number F-1756 to L.M.C.); a Fulbright Garcia-Robles Fellowship [to A.M.R.N]; the National Institute of Health [project number 5R01GM135495-04 to L.M.C., T.R.S., and A.M.R.N.]; and a University of Texas at Austin Continuing Graduate Fellowship [to A.M.R.N]. Figures and diagrams were prepared in BioRender and are accessible at the following links: Figure 1 (BioRender.com/o05u602), Figure 3, (BioRender.com/s22k096), Figure 4

(BioRender.com/g88x379), Figure 5 (BioRender.com/c79o110), Figure 6 (), and the Graphical Abstract (BioRender.com/f14x732). Data were visualized using GraphPad Prism.

