## Supplementary Information for "Concentration dependent CsrA regulation of the *uxuB* transcript leads to development of post-transcriptional BANDPASS Filter"

### Table of Contents

**Supplementary Materials and Methods**

**Supplementary Tables**
Table S1 – Strains and Plasmids used in this study
Table S2 – Oligonucleotides used in this study

**Supplementary Figures**
Figure S1 – EMSA Screen for all putative mRNA targets identified to interact with CsrA through TriFC Assay
Figure S2 – Mutational analysis for the *uxuB* leader sequence by EMSA
Figure S3 – *In vivo* reporter assay for 5' UTR and CDS Mutant *uxuB* leader sequences
Figure S4 – The CsrA-*uxuB* interaction does not significantly impact utilization of hexuronic acids
Figure S5 – Benchmarking the Csr-regulated BANDPASS Filter against a transcriptionally regulated BANDPASS filter
Figure S6 – Time course of BANDPASS Filter fold activation at different induction levels

**Supplementary References**

### Supplementary Materials and Methods.

#### Protein purification

His-tagged CsrA (CsrA-H6) was purified via Ni-NTA affinity chromatography as previously described in (Rojano-Nisimura, Simmons, et al., 2023). Briefly, a pET-21a (+) vector (pCSB12) was used to express CsrA-H6 upon IPTG induction in *E. coli* BL21 (DE3). Protein expression was allowed to proceed for a total of ~20 hrs. Afterwards, CsrA-H6 was recovered from the soluble fraction (~10 mL) via nickel column purification using a Ni-NTA agarose column resin (Qiagen). Bound CsrA was recovered in a 50% elution buffer (250 mM imidazole) and the recovered fractions were analyzed by SDS-PAGE to confirm CsrA expression. Fractions containing recovered CsrA were exchanged into CsrA storage buffer (10 mM Tris-HCl, 100 mM KCl, 10 mM $MgCl_2$, 25% glycerol, pH 7.0). Protein concentration was determined by Bradford assay and protein purity was validated by LC-MS/MS. Protein aliquots were stored in single-use aliquots at -20°C.

#### Growth assays

Overnight cultures of *E. coli* MG166 and E. coli MG1655 uxuB:GGA (which has its genomic copy of the *uxuB* gene substituted by that of the "uxuB_no_GGAs" mutant; see Materials and Methods for a description of the protocol used to generate genomic mutants) were grown overnight in 5 mL cultures of LB media. The next day, new cultures were seeded 1:100 in 5 mL M9 Media supplemented with 0.2% Glucose and grown overnight. These saturated cultures were again diluted 1:100 into either M9 + 0.2% Glucose, M9 + 0.2% Glucuronate, or M9 + 0.1% Glucose + 0.1% Glucuronate. These cultures were seeded in 200 uL in honeycomb plates (Growth Curves USA) and incubated in a BioScreen C Type FP-1100-C Analysis System (Thermo Labsystems). $OD_{600}$ was measured every 15 minutes for 14 hours until cells reached stationary phase. The growth rates were analyzed using the *growthrates* package in R.

### Supplementary Tables

**Supplementary Table S1.** Stains and Plasmids used in this study

| Strain or Plasmid | Description/Genotype | Reference |
|---|---|---|
| *E. coli DH5α* | *fhuA2 lac(del)U169 phoA glnV44 Φ80' lacZ(del)M15 gyrA96 recA1 relA1 endA1 thi-1 hsdR17* | Contreras Lab, U. of Texas at Austin |
| *E. coli* MG1655 | *F- lambda- ilvG- rfb-50 rph-1* | Contreras Lab, U. of Texas at Austin |
| *E. coli* BL21 (DE3) | *fhuA2 [lon] ompT gal (λ DE3) [dcm] ∆hsdS λ DE3 = λ sBamHIo ∆EcoRI-B int::(lacI::PlacUV5::T7 gene1) i21 ∆nin5* | New England Biolabs |
| *ΔcsrB* | *MG1655 ΔcsrB ΔglgCAP ΔpgaABCD lacI::lacI$^q$* | (1) |
| *Δcsr* | *MG1655 ΔcsrA::cam ΔcsrB ΔcsrC ΔcsrD ΔglgCAP ΔpgaABCD ΔlacI::lacI$^q$* | (2) |
| *uxuB::uxuB GGAs mutant (uxuB_no_GGAs)* | *MG1655 ΔuxuB::uxuB -40C>G -35C>G -16G>C -9G>C +47 G>T +48G>T +64G>C +84C>A* | This work |
| pTriFC-astD | astD 5'UTR- MS2BD-rrnB fusion. SpeI/SphI template digestion. astD insert amplified by colony PCR. Gibson Assembly. | This work |
| pTriFC-cmk | cmk 5'UTR- MS2BD-rrnB fusion. SpeI/SphI template digestion. cmk insert amplified by colony PCR. Gibson Assembly. | This work |
| pTriFC-entC | entC 5'UTR- MS2BD-rrnB fusion. SpeI/SphI template digestion. entC insert amplified by colony PCR. Gibson Assembly. | This work |
| pTriFC-pepT | pepT 5'UTR- MS2BD-rrnB fusion. SpeI/SphI template digestion. pepT insert amplified by colony PCR. Gibson Assembly. | This work |
| pTriFC-rodZ | rodZ 5'UTR- MS2BD-rrnB fusion. SpeI/SphI template digestion. rodZ insert amplified by colony PCR. Gibson Assembly. | This work |
| pTriFC-truC | truC 5'UTR- MS2BD-rrnB fusion. SpeI/SphI template digestion. truC insert amplified by colony PCR. Gibson Assembly. | This work |
| pTriFC-uxuB | uxuB 5'UTR- MS2BD-rrnB fusion. SpeI/SphI template digestion. uxuB insert amplified by colony PCR. Gibson Assembly. | This work |
| pTriFC-yqjE | yqjE 5'UTR- MS2BD-rrnB fusion. SpeI/SphI template digestion. yqjE insert amplified by colony PCR. Gibson Assembly. | This work |
| pTriFC-bfr | bfr 5'UTR- MS2BD-rrnB fusion. | (3) |
| pTriFC-dps | dps 5'UTR- MS2BD-rrnB fusion. | (3) |
| pTriFC-fecA | fecA 5'UTR- MS2BD-rrnB fusion. | (3) |
| pTriFC-patA | patA 5'UTR- MS2BD-rrnB fusion. | (3) |
| pTriFC-purM | purM 5'UTR- MS2BD-rrnB fusion. | (3) |
| pTriFC-ybaL | ybaL 5'UTR- MS2BD-rrnB fusion. | (3) |
| pTriFC-glgC | glgC 5'UTR- MS2BD-rrnB fusion. | (3) |
| pMS2-CYFP | MS2-linker-CYFP fusion | (4) |

| | | |
|---|---|---|
| pCSB12 | *csrA* insert at the NdeI and BamHI sites of pET21a+ vector. | (5) |
| pHL600 | pZ with p15a origin of replication. *csrA* is expressed from a pLlacO-1 promoter. | (2) |
| pHL1756-phoB | *phoB* 5'UTR- *gfp* translational fusion. SalI/SphI template digestion. *phoB* insert amplified by colony PCR. Gibson Assembly. | (6) |
| pHL1756-gmk | *gmk* 5'UTR- *gfp* translational fusion. SalI/SphI template digestion. *gmk* insert amplified by colony PCR. Gibson Assembly. | (3) |
| pHL1756-fecA | *fecA* 5'UTR- *gfp* translational fusion. SalI/SphI template digestion. *fecA* insert amplified by colony PCR. Gibson Assembly. | (3) |
| pHL1756-glgC | *glgC* 5'UTR- *gfp* translational fusion. SalI/SphI template digestion. *glgC* insert amplified by colony PCR. Gibson Assembly. | (3) |
| pHL1756-uxuB | *uxuB* 5'UTR- *gfp* translational fusion. SalI/SphI template digestion. uxuB insert amplified by colony PCR. Gibson Assembly. | (3) |
| pHL1756-uxuB-UPSTR-mut | *uxuB-UP* 5'UTR- *gfp* translational fusion. SalI/SphI template digestion. Insert synthesized as gBlock. Gibson Assembly. | This work |
| pHL1756-uxuB-DOWNSTR-mut | *uxuB-DOWN* 5'UTR- *gfp* translational fusion. SalI/SphI template digestion. Insert synthesized as gBlock. Gibson Assembly. | This work |
| pHL1756-uxuB-no GGAs-mut | *uxuB-noGGAs* 5'UTR- *gfp* translational fusion. SalI/SphI template digestion. Insert synthesized as gBlock. Gibson Assembly. | This work |
| pHL1756-uxuB-mut-2 | *uxuB-mut2* 5'UTR- *gfp* translational fusion. SalI/SphI template digestion. Insert synthesized as gBlock. Gibson Assembly. | This work |
| pBP-uxuB-GFP-WT CsrB | BANDPASS Filter Plasmid: *uxuB 5'UTR-gfpmut3* ($P_{con12}$), WT CsrB ($P_{LlacO}$: lacI), ColE1 ori, CarbR | This work |
| pBP-uxuB-GFP-H11 CsrB | BANDPASS Filter Plasmid: *uxuB 5'UTR-gfpmut3* ($P_{con12}$), H11 CsrB ($P_{LlacO}$: lacI), ColE1 ori, CarbR | This work |
| pBP-uxuB-GFP-L2 CsrB | BANDPASS Filter Plasmid: *uxuB 5'UTR-gfpmut3* ($P_{con12}$), L2 CsrB ($P_{LlacO}$: lacI), ColE1 ori, CarbR | This work |
| pBP-GFP-DAS CsrB-WT | BANDPASS Filter Plasmid: *uxuB 5'UTR-gfpmut3*-DAS Degradation Tag ($P_{con12}$), WT CsrB ($P_{LlacO}$: lacI), ColE1 ori, CarbR | This work |
| pTRS063 | cBUFFER-mCherry: *glgC 5' UTR*-TTGGT-*mcherry* ($P_{con}$12), WT CsrB ($P_{LlacO}$; lacI), p15a ori, KanR | (7) |
| pDIM-C8-TEM1 | *Bla-TEM1* ($P_{tac}$: lacI$^q$), p15a ori, CmR | (8) |
| pTS01 | ampR ($P_{ampP/O}$), *gfpmut3-tetC* ($P_{ampP/O}$), *lacI$^q$*, CloDF13 ori, SmR | (8) |

**Supplementary Table S2.** Oligonucleotides used in this study.

| Primer Name | Sequence | Purpose |
|---|---|---|
| astD FWD | gctttaaatttgcgcacactagtCCGAGCGTTTGATTTTAAC | pTriFC cloning |
| astD RV | cacacgtcttcgcatgcCATCATTGCCTTGCCATAAC | pTriFC cloning |
| cmk FWD | gctttaaatttgcgcacactagtTTATGTTAACGGTACGCCTGTTTTAAG | pTriFC cloning |
| cmk RV | cacacgtcttcgcatgcGCAGATGCCATTGCAACG | pTriFC cloning |
| entC FWD | gctttaaatttgcgcacactagtGAAAATATAAATGATAATCATTATTAAAG | pTriFC cloning |
| entC RV | cacacgtcttcgcatgcCTGACGTCGTAAAACTGC | pTriFC cloning |
| pepT FWD | gctttaaatttgcgcacactagtACAAAAAGTGAGGGTGACTACATG | pTriFC cloning |
| pepT RV | cacacgtcttcgcatgcACTTCCATTGGCCTTCCG | pTriFC cloning |
| rodZ FWD | gctttaaatttgcgcacactagtAACGGTTAACTTAACGGATGTTTC | pTriFC cloning |
| rodZ RV | cacacgtcttcgcatgcCAACGGCCTGCTGACTAAG | pTriFC cloning |
| truC FWD | gctttaaatttgcgcacactagtCCTATTATGAAATGGCGC | pTriFC cloning |
| truC RV | cacacgtcttcgcatgcCTACTTTCTCGTCGCGATC | pTriFC cloning |
| uxuB FWD | gctttaaatttgcgcacactagtGCGCTTTCTTTAGCCGTTAATATCCAC | pTriFC cloning |
| uxuB RV | cacacgtcttcgcatgcGAAACGCCCCGCAACCGA | pTriFC cloning |
| yqjE FWD | gctttaaatttgcgcacactagtCGCGTGCCGATGAGTATGTGC | pTriFC cloning |
| yqjE RV | cacacgtcttcgcatgcTCGCTGCCCGATGCCCAG | pTriFC cloning |
| entC-fwd-T7 | GATCTAATACGACTCACTATAGGGAGgtttgcttttaggttagcgaccGA | IVT for EMSAs |
| yqjE-fwd-T7 | GATCTAATACGACTCACTATAGGGAGCGCGTGCCGATG AGTATGTGCGCGAAAATC | IVT for EMSAs |
| cmk-fwd-T7 | GATCTAATACGACTCACTATAGGGAGTTATGTTAACGGTACGC CTGTTTTAAGGAG | IVT for EMSAs |
| dps-fwd-T7 | GATCTAATACGACTCACTATAGGGAGGTTAATTACTGGGACAT AACATCAAGAGG | IVT for EMSAs |
| ybaL-fwd-T7 | GATCTAATACGACTCACTATAGGGAGGTAATTTTGGTTTTCCC GGCCAAAAATGG | IVT for EMSAs |
| rodZ-fwd-T7 | GATCTAATACGACTCACTATAGGGAGAACGGTTAACTTAACGG ATGTTTCGCGGTG | IVT for EMSAs |
| uxuB-fwd-T7 | GATCTAATACGACTCACTATAGGGAGGCGCTTTCTTTAGCCGT TAATATCCACCGG | IVT for EMSAs |
| aidB-fwd-T7 | GATCTAATACGACTCACTATAGGGAGGTGACTGCCATTGATGG AGGGAGACAC | IVT for EMSAs |
| patA-fwd-T7 | GATCTAATACGACTCACTATAGGGAGATTTGTGTTTATCCCGAT TTTCGCGATCGC | IVT for EMSAs |
| fecA-fwd-T7 | GATCTAATACGACTCACTATAGGGAGTTCTCGTTCGACTCATA GCTGAACACAACA | IVT for EMSAs |
| phoB-fwd-T7 | GATCTAATACGACTCACTATAGGGAGGCATTAATGATCGCAAC CTATTT | IVT for EMSAs |
| entC-rev-T7 | CTGACGTCGTAAAACTGCGGTAC | IVT for EMSAs |
| yqjE-rev-T7 | TCGCTGCCCGATGCCCAGAACGCTTTTA | IVT for EMSAs |
| cmk-rev-T7 | GCAGATGCCATTGCAACGCTT | IVT for EMSAs |
| dps-rev-T7 | TCAGCAACTCTACTGTTGCTTTTTTC | IVT for EMSAs |
| ybaL-rev-T7 | CCAGAGGAGAAATACGTAGTTTATTGGCCA | IVT for EMSAs |

| | | |
|---|---|---|
| rodZ-rev-T7 | CAACGGCCTGCTGACTAAGTCCTAGTTGTT | IVT for EMSAs |
| uxuB-rev-T7 | GAAACGCCCCGCAACCGAGAT | IVT for EMSAs |
| aidB-rev-T7 | CACGCGTTACCGCTTCGCAGA | IVT for EMSAs |
| patA-rev-T7 | GTGCTTTCATCTCCTCATGATCCAGCG | IVT for EMSAs |
| fecA-rev-T7 | CAGCAAAAGCGGAAAACGAGAGACC | IVT for EMSAs |
| phoB-rev-T7 | cataatcttccgcttcgacc | IVT for EMSAs |
| uxuAB_UTR_probe_7_FWD | AATTCAGAATCCGTCTTCAAAT | Cloning CF-iRS$^3$ plasmids |
| uxuAB_UTR_probe_7_RVS | TGGTATTTGAAGACGGATTCTG | Cloning CF-iRS$^3$ plasmids |
| uxuAB_UTR_probe_8_FWD | AATTCCATCGTAAACTCCAGAT | Cloning CF-iRS$^3$ plasmids |
| uxuAB_UTR_probe_8_RVS | TGGTATCTGGAGTTTACGATGG | Cloning CF-iRS$^3$ plasmids |
| uxuAB_UTR_probe_15_FWD | AATTCATCCCATGACGGGCGGT | Cloning CF-iRS$^3$ plasmids |
| uxuAB_UTR_probe_15_RVS | TGGTACCGCCCGTCATGGGATG | Cloning CF-iRS$^3$ plasmids |
| uxuAB_UTR_probe_16_FWD | AATTCATTCCAGACGAGAATGT | Cloning CF-iRS$^3$ plasmids |
| uxuAB_UTR_probe_16_RVS | TGGTACATTCTCGTCTGGAATG | Cloning CF-iRS$^3$ plasmids |
| uxuAB_UTR_probe_19_FWD | AATTCCGAGATGCACAATT | Cloning CF-iRS$^3$ plasmids |
| uxuAB_UTR_probe_19_RVS | TGGTAATTGTGCATCTCGG | Cloning CF-iRS$^3$ plasmids |
| T7-uxuB-wild type | TAATACGACTCACTATAGGGATAATGCGCTTTCTTTAGCCGTTAATATCCACCGGCATGGCTGCGCGCCGTGCCGGTTCCTTCTTCCTTGCCGTCACTCTTTGAAGACGGATTCTGGAGTTTACGATGACTACTATTGTTGACAGCAATCTGCCGGTTGCCCGCCCGTCATGGGATCATTCTCGTCTGGAATCACGCATTGTGCATCTCGGTTGCGGGGCGTTTC | EMSAs |
| T7-uxuB-UPSTR mut | TAATACGACTCACTATAGGGATAATGCGCTTTCTTTAGCCGTTAATATCCACCGGCATGGCTGCGCGCCGTGCCGGTTCCTTCTTgCTTGgCGTCACTCTTTGAAGACGcATTCTGcAGTTTACGATGAC | EMSAs |

| | | |
|---|---|---|
| | TACTATTGTTGACAGCAATCTGCCGGTTGCCCGCCCGTCATGG GATCATTCTCGTCTGGAATCACGCATTGTGCATCTCGGTTGCG GGGCGTTTCGCAtGC | |
| T7-uxuB-DOWNSTR mut | TAATACGACTCACTATAGGGATAATGCGCTTTCTTTAGCCGTTA ATATCCACCGGCATGGCTGCGCGCCGTGCCGGTTCCTTCTTC CTTGCCGTCACTCTTTGAAGACGGATTCTGGAGTTTACGATGA CTACTATTGTTGACAGCAATCTGCCGGTTGCCCGCCCGTCATtt GATCATTCTCGTCTGcAATCACGCATTGTGCATCTaGGTTGCG GGGCGTTTCGCAtGC | EMSAs |
| T7-uxuB-no GGAs mut | TAATACGACTCACTATAGGGATAATGCGCTTTCTTTAGCCGTTA ATATCCACCGGCATGGCTGCGCGCCGTGCCGGTTCCTTCTTg CTTGgCGTCACTCTTTGAAGACGcATTCTGcAGTTTACGATGAC TACTATTGTTGACAGCAATCTGCCGGTTGCCCGCCCGTCATttG ATCATTCTCGTCTGcAATCACGCATTGTGCATCTaGGTTGCGG GGCGTTTCGCAtGC | EMSAs |
| T7-uxuB-UPSTR-trunc-wild type | TAATACGACTCACTATAGGGATAATGCGCTTTCTTTAGCCGTTA ATATCCACCGGCATGGCTGCGCGCCGTGCCGGTTCCTTCTTC CTTGCCGTCACTCTTTGAAGACGGATTCTGGAGTTTACGA | EMSAs |
| T7-uxuB-UPSTR-trunc-mut | TAATACGACTCACTATAGGGATAATGCGCTTTCTTTAGCCGTTA ATATCCACCGGCATGGCTGCGCGCCGTGCCGGTTCCTTCTTg CTTGgCGTCACTCTTTGAAGACGcATTCTGcAGTTTACGA | EMSAs |
| T7-uxuB-DOWNSTR-trunc-wild type | TAATACGACTCACTATAGGGATAATATGACTACTATTGTTGACA GCAATCTGCCGGTTGCCCGCCCGTCATGGGATCATTCTCGTC TGGAATCACGCATTGTGCATCTCGGTTGCGGGGCGTTTC | EMSAs |
| T7-uxuB-DOWNSTR-trunc-mut | TAATACGACTCACTATAGGGATAATATGACTACTATTGTTGACA GCAATCTGCCGGTTGCCCGCCCGTCATttGATCATTCTCGTCTG cAATCACGCATTGTGCATCTaGGTTGCGGGGCGTTTC | EMSAs |
| uxuB_fwd-IVTT | tcactataggtaccggtGCGCTTTCTTTAGCCGTTAATATCC | IVTTs |
| uxuB_rev-IVTT | ttcttcacctttgctcatGAAACGCCCCGCAACCGAG | IVTTs |
| uxuB-mut-2 | gctttaaatttgcgcacactagtGCGCTTTCTTTAGCCGTTAATATCCACC GGCATGGCTGCGCGCCGTGCCGGTTCCTTCTTCCTTGCCGTC ACTCTTTGAAGACGGATTCTGGAGTTTACaATGACTACTATTGT TGACAGCAATCTGCCGGTTGCCCGCCCGTCATGGGATCATTC TCGTCTGGAATCACGCATTGTGCATCTCGGTTGCGGGGCGTT TCgcatgcgaagacgtgtg | gBlock for reporter cloning |
| uxuB-UP-mut | gtggattccacacaggtcgacGCGCTTTCTTTAGCCGTTAATATCCACC GGCATGGCTGCGCGCCGTGCCGGTTCCTTCTTgCTTGgCGTC ACTCTTTGAAGACGcATTCTGcAGTTTACGATGACTACTATTGT TGACAGCAATCTGCCGGTTGCCCGCCCGTCATGGGATCATTC TCGTCTGGAATCACGCATTGTGCATCTCGGTTGCGGGGCGTT TCGCAtGCgtaaaggagaag | gBlock for reporter cloning |
| uxuB-DOWN-mut | gtggattccacacaggtcgacGCGCTTTCTTTAGCCGTTAATATCCACC GGCATGGCTGCGCGCCGTGCCGGTTCCTTCTTCCTTGCCGTC ACTCTTTGAAGACGGATTCTGGAGTTTACGATGACTACTATTGT TGACAGCAATCTGCCGGTTGCCCGCCCGTCATttGATCATTCTC GTCTGcAATCACGCATTGTGCATCTaGGTTGCGGGGCGTTTCG CAtGCgtaaaggagaag | gBlock for reporter cloning |

| | | |
|---|---|---|
| uxuB-noGGA-UTR-OH | gtggattccacacaggtcgacGCGCTTTCTTTAGCCGTTAATATCCACC<br>GGCATGGCTGCGCGCCGTGCCGGTTCCTTCTTgCTTGgCGTC<br>ACTCTTTGAAGACGcATTCTGcAGTTTACGATGACTACTATTGT<br>TGACAGCAATCTGCCGGTTGCCCGCCCGTCATttGATCATTCTC<br>GTCTGcAATCACGCATTGTGCATCTaGGTTGCGGGGCGTTTCG<br>CAtGCgtaaaggagaag | gBlock for reporter cloning |
| rTRS328 | gcaataactagcataaccc | FW Primer to amplify pCDF |
| rTRS329 | atggtatatctccttattaaagttaa | RV Primer to amplify pCDF |
| rTRS330 | ACCGCTGAGCAATAACTAGC | FW Primer to amplify pAcyc |
| rTRS331 | GCGCAACGCAATTAATGTAAGT | RV Primer to amplify pAcyc |
| rTRS332 | atgagtaaaggagaagaacttttc | FW Primer to amplify GFPmut3 from start |
| rTRS337 | agaagcgtcagcgtagttttcgtcgttagcagctttgtagagctcatccatgcca | FW Gibson Primer to amplify towards gfpmut3 with DAS tag on bandpass plasmids |
| rTRS338 | gctgctaacgacgaaaactacgctgacgcttcttaacttggctgttttggcgg | RV Gibson Primer to amplify towards rrnB12 terminator with DAS tag on bandpass plasmids |
| TS1-gblk1 | ttaactttaataaggagatataccatttaTTTGTGCAGCACCCCGGTCAACCA<br>ACGGGAAAATTCACGCATCGCGGGCGTCTCCGGGCGAGATTG<br>CAAACGCGTTATCCAGTAGCTCCCCAAATCAATCTGCGTTAAA<br>AACGGCTGAACGATACGTTCACTGCTGAGTAAATGCGTGAACA<br>TTCTGACTGGCGCAATTGCCACACCGCTCCCCGCCTGAGCGG<br>CTTCCAGCATGGTGACGGACGAATCAAACACCATCACATTGTG<br>CGTCGGTGACGGAGGCGCCTCTCCGGCCGCCTGCATCCAAA<br>GCGCCCATTCATCCCGCCGATACGATCGTAATAACGGAAATTT<br>CAGGATATCCGCAGGCGTTTGAATCTGCGAAGCCAATGTTGG<br>CGAACACAGTGGAGACATCAGCGCGCTACATAAATATTGCGC<br>ATCGGTATCGTGCCAAGCTCCCCCACCGTAGCGAATGGTATA<br>ATCCAGCCCTTCGGCGGCGGGATCCACGCGATTATTATGGGT<br>AGAAATATGCAAATCAATATGTGGGTAACTGCGCTTAAAATCG<br>CTCAACAGCGGGAAAAGACAGCCGATAGCAAAGGTTCCCACT<br>ACGCCAATTTTAAGTTTTTCCTGGGTCTGTTTAGTGGCAAAAC<br>GATCCAACATCCCCGCCATACGATCGAAGGAGTCATTAAGTAC<br>AGGTAACAGACTTTCCCCTTCAGTCGTCAACATTAATCCACGA<br>GAACCGCGCACAAAAAGCTGACAATTCAGCTGCTGCTCCAGC<br>GATTTGACATGCTGGCTGATGGCAGAATGCGTCACGTTGAGC<br>TCAATCGCAGCGCGGGTAAAGCTGAGATGTCTGGCCGCGGCT<br>TCAAAAGCCCGCAGCGAGTTAAGAGGGATATAGCTACGCGTca<br>tCATTAAGCCTGTTAGAAAAACTTATATCTGCTGCTAAATTTAAC<br>CGTTTGTCAACACGGTGCAAATCAAACACACTGATTGCGTCTG | gblock to insert Band pass gate |

| | | |
|---|---|---|
| | ACGGGCCCGGACACCCTTTTGTTTTTAATTACGGAACTGATTT<br>CATGATGAGTAAAGGAGAAGAACTTTTCACTGGAGTTGTCCCA<br>ATTCTTGTTGAATTAGATGGTGATGTTAATGGGCACAAATTTTC<br>TGTCAGTGGAGAGGGTGAAGGTGATGCAACATACGGAAAACT<br>TACCCTTAAATTTATTTGCACTACTGGAAAACTACCTGTTCCAT<br>GGCCAACACTTGTCACTACTTTCGGTTATGGTGTTCAATGCTTT<br>GCGAGATACCCAGATCATATGAAACAGCATGACTTTTTCAAGA<br>GTGCCATGCCCGAAGGTTATGTACAGGAAAGAACTATATTTTT<br>CAAAGATGACGGGAACTACAAGACACGTGCTGAAGTCAAGTTT<br>GAAGGTGATACCCTTGTTAATAGAATCGAGTTAAAAGGTATTG<br>ATTTTAAAGAAGATGGAAACATTCTTGGACACAAATTGGAATAC<br>AACTATAACTCACACAATGTATACATCATGGCAGACAAACAAAA<br>GAATGGAATCAAAGTTAACTTCAAAATTAGACAC | |
| TS1-gblk2 | GGAATCAAAGTTAACTTCAAAATTAGACACAACATTGAAGATG<br>GAAGCGTTCAACTAGCAGACCATTATCAACAAAATACTCCAATT<br>GGCGATGGCCCTGTCCTTTTACCAGACAACCATTACCTGTCCA<br>CACAATCTGCCCTTTCGAAAGATCCCAACGAAAAGAGAGACCA<br>CATGGTCCTTCTTGAGTTTGTAACAGCTGCTGGGATTACACAT<br>GGCATGGATGATCTCTACAAATAAatgaaatctaacaatgcgctcatcgtcat<br>cctcggcaccgtcaccctggatgctgtaggcataggcttggttatgccggtactgccgggcct<br>cttgcgggatatcgtccattccgacagcatcgccagtcactatggcgtgctgctagcgctatat<br>gcgttgatgcaatttctatgcgcacccgttctcggagcactgtccgaccgctttggccgccgcc<br>cagtcctgctcgcttcgctacttggagccactatcgactacgcgatcatggcgaccacaccc<br>gtcctgtggatcctctacgccggacgcatcgtggccggcatcaccggcgccacaggtgcgg<br>ttgctggcgcctatatcgccgacatcaccgatggggaagatcgggctcgccacttcgggctc<br>atgagcgcttgtttcggcgtgggtatggtggcaggccccgtggccgggggactgttgggcgc<br>catctccttgcatgcaccattccttgcggcggcggtgctcaacggcctcaacctactactggg<br>ctgcttcctaatgcaggagtcgcataagggagagcgtcgaccgatgcccttgagagccttca<br>acccagtcagctccttccggtgggcgcggggcatgactatcgtcgccgcacttatgactgtctt<br>ctttatcatgcaactcgtaggacaggtgccggcagcgctctgggtcattttcggcgaggaccg<br>ctttcgctggagcgcgacgatgatcggcctgtcgcttgcggtattcggaatcttgcacgccctc<br>gctcaagccttcgtcactggtcccgccaccaaacgtttcggcgagaagcaggccattatcgc<br>cggcatggcggccgacgcgctgggctacgtcttgctggcgttcgcgacgcgaggctggatg<br>gccttccccattatgattcttctcgcttccggcggcatcgggatgcccgcgttgcaggccatgct<br>gtccaggcaggtagatgacgaccatcagggacagcttcaaggatcgctcgcggctcttacc<br>agcctaacttcgatcactggaccgctgatcgtcacggcgatttatgccgcctcggcgagcac<br>atggaacgggttggcatggattgtaggcgccgccctataccttgtctgcctccccgcgttgcgt<br>cgcggtgcatggagccgggccacctcgacctgagcaataactagcataacccctt | gblock to insert<br>Band pass gate |
| TEM1-<br>gblck | ACTTACATTAATTGCGTTGCGCGTTGACAATTAATCATCGGCTC<br>GTATAATGTGTGGAATTGTGAGCGGATAACAATTTCACACAGA<br>ACAGAATTCTAATGAGTATTCAACATTTCCGTGTCGCCCTTATT<br>CCCTTTTTTGCGGCATTTTGCCTTCCTGTTTTTGCTCACCCAGA<br>AACGCTGGTGAAAGTAAAAGATGCTGAAGATCAGTTGGGTGC<br>ACGAGTGGGTTACATCGAACTGGATCTCAACAGCGGTAAGAT<br>CCTTGAGAGTTTTCGCCCCGAAGAACGTTTTCCAATGATGAGC<br>ACTTTTAAAGTTCTGCTATGTGGCGCGGTATTATCCCGTGTTG<br>ACGCCGGGCAAGAGCAACTCGGTCGCCGCATACACTATTCTC<br>AGAATGACTTGGTTGAGTACTCACCAGTCACAGAAAAGCATCT<br>TACGGATGGCATGACAGTAAGAGAATTATGCAGTGCTGCCATA<br>ACCATGAGTGATAACACTGCGGCCAACTTACTTCTGACAACGA<br>TCGGAGGACCGAAGGAGCTAACCGCTTTTTTGCACAACATGG<br>GGGATCATGTAACTCGCCTTGATCGTTGGGAACCGGAGCTGA | gblock to insert<br>Ptac-TEM1 CDS<br>into pAcyc |

| | |
|---|---|
| | ATGAAGCCATACCAAACGACGAGCGTGACACCACGATGCCTG CAGCAATGGCAACAACGTTGCGCAAACTATTAACTGGCGAACT ACTTACTCTAGCTTCCCGGCAACAATTAATAGACTGGATGGAG GCGGATAAAGTTGCAGGACCACTTCTGCGCTCGGCCCTTCCG GCTGGCTGGTTTATTGCTGATAAATCTGGAGCCGGTGAGCGT GGGTCTCGCGGTATCATTGCAGCACTGGGGCCAGATGGTAAG CCCTCCCGTATCGTAGTTATCTACACGACGGGGAGTCAGGCA ACTATGGATGAACGAAATAGACAGATCGCTGAGATAGGTGCCT CACTGATTAAGCATTGGTAAACCGCTGAGCAATAACTAGCA |

### Supplementary Figures.

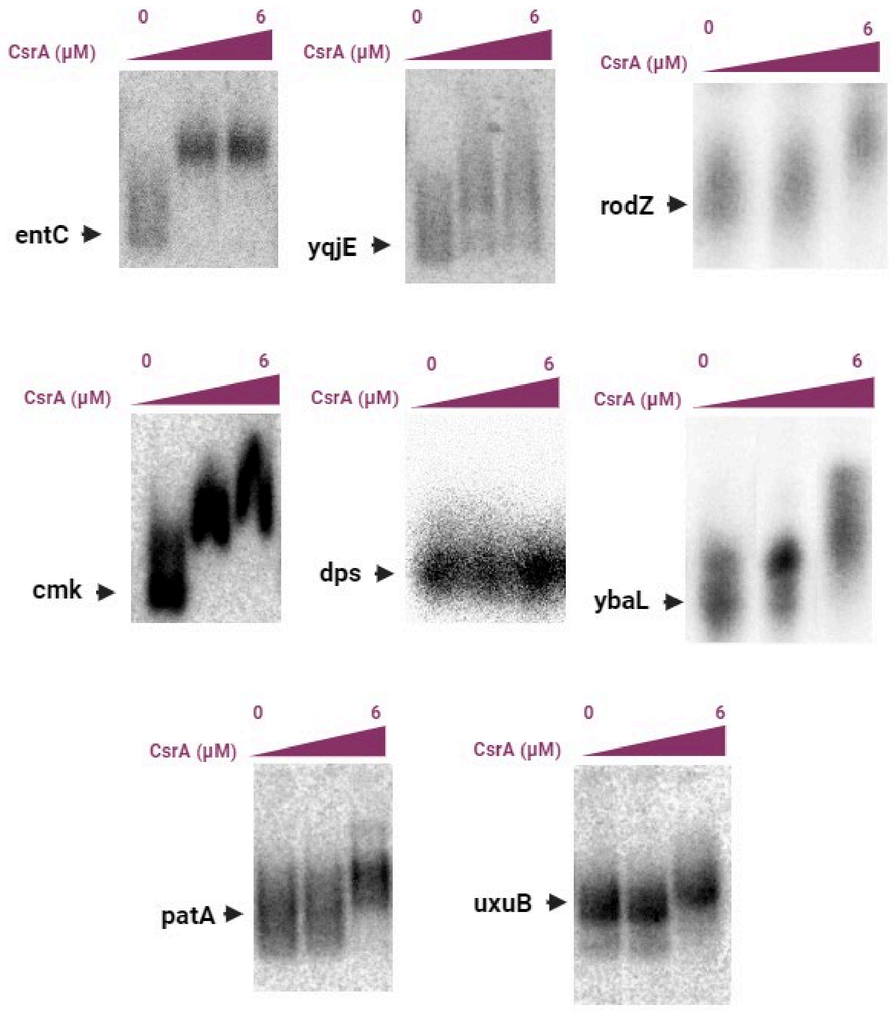

**Supplementary Figure S1. EMSA Screen for all putative mRNA targets identified to interact with CsrA through TriFC Assay.** Electrophoretic mobility shift assays (EMSAs) were performed for each of the target mRNA candidates identified from the TriFC assay to evaluate *in vitro* CsrA-RNA binding. For screening, 10 nM of radiolabeled RNAs were incubated with 0, 3 (300:1 ratio) or 6 (600:1 ratio) μM of purified CsrA**.**

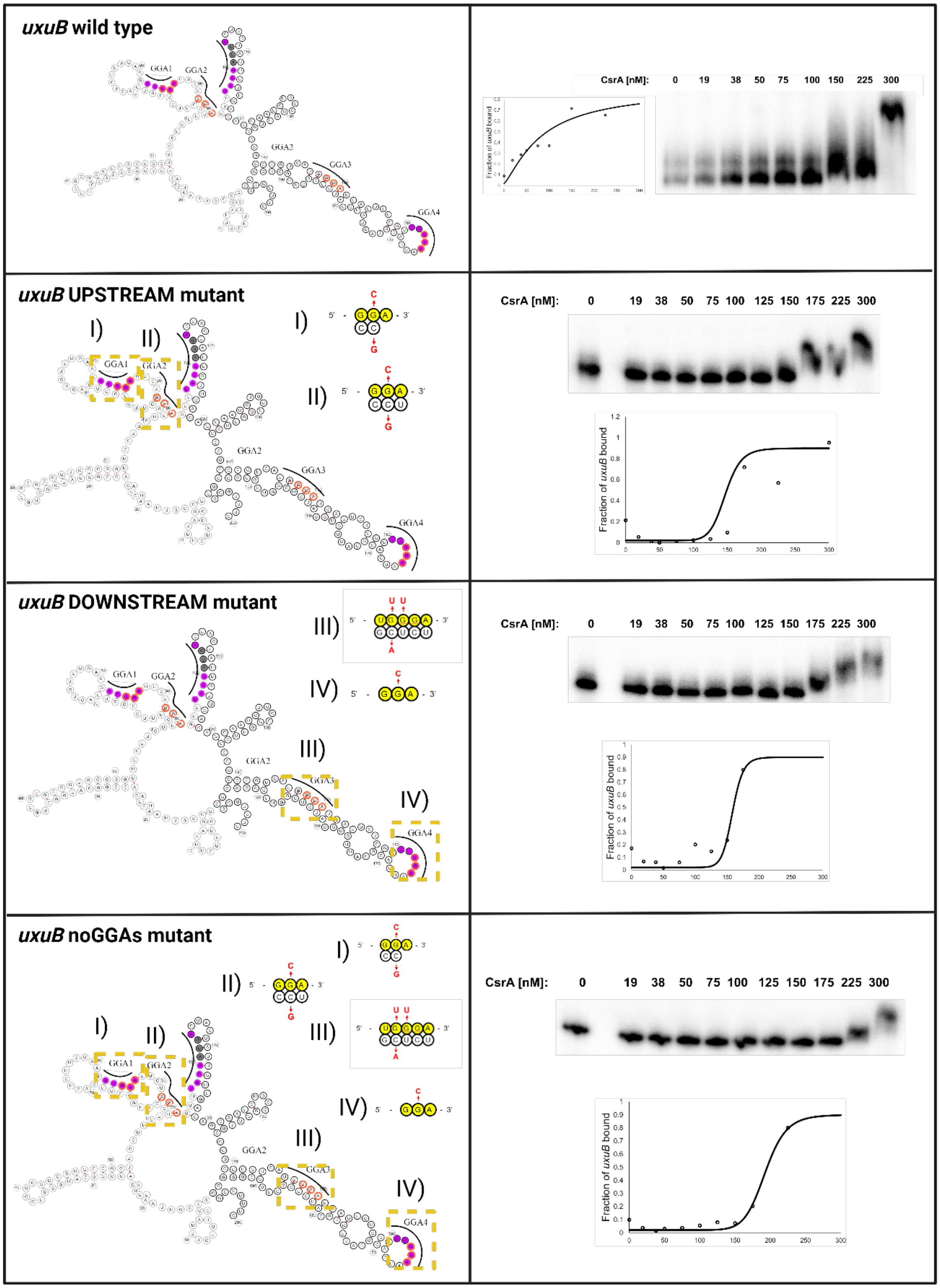

(Continued on next page).

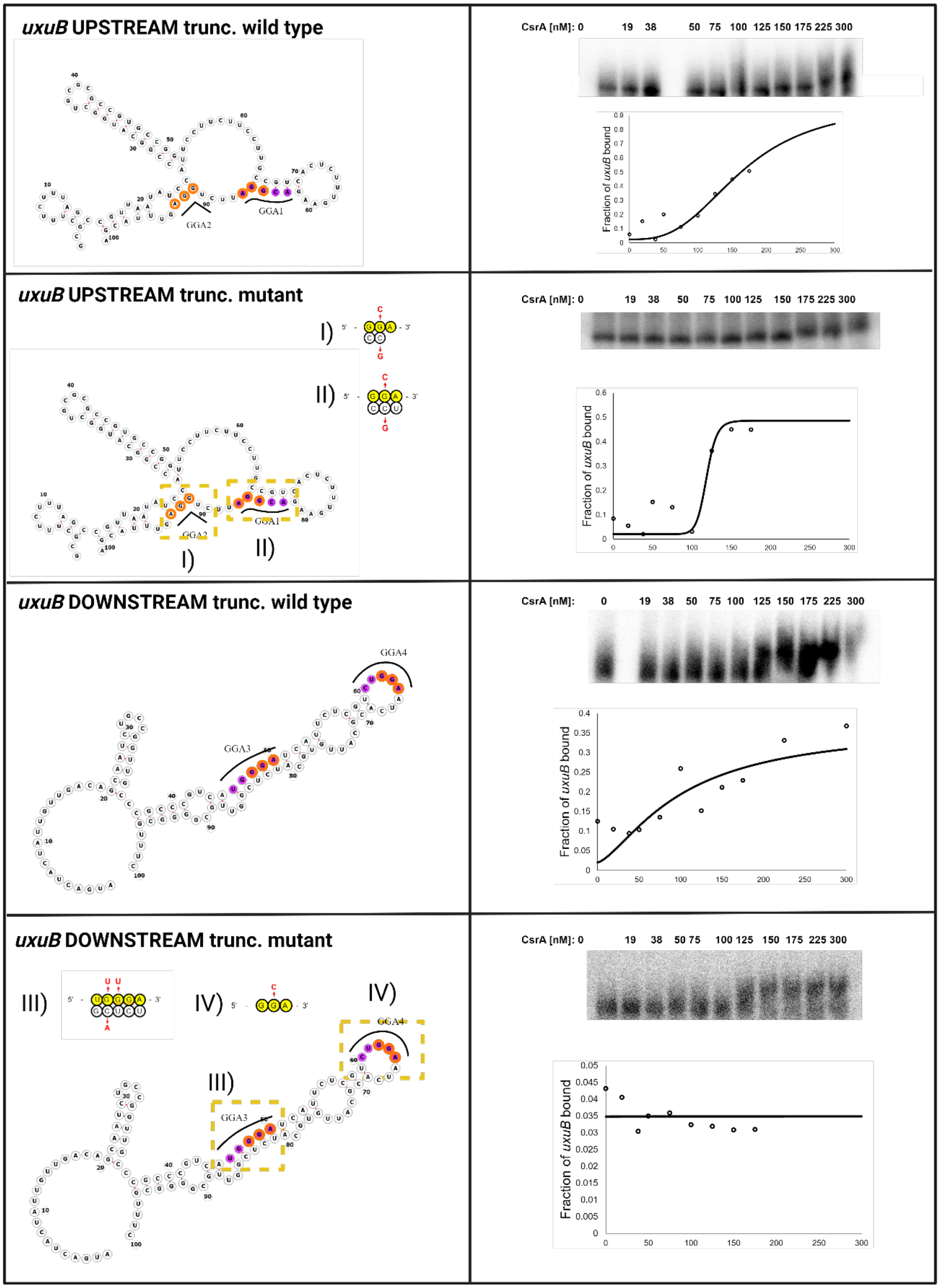

uxuB UPSTREAM trunc. wild type
GGA1
GGA2
CsrA [nM]: 0 19 38 50 75 100 125 150 175 225 300
Fraction of uxuB bound
uxuB UPSTREAM trunc. mutant
I)
II)
GGA1
GGA2
CsrA [nM]: 0 19 38 50 75 100 125 150 175 225 300
Fraction of uxuB bound
uxuB DOWNSTREAM trunc. wild type
GGA3
GGA4
CsrA [nM]: 0 19 38 50 75 100 125 150 175 225 300
Fraction of uxuB bound
uxuB DOWNSTREAM trunc. mutant
III)
IV)
GGA3
GGA4
CsrA [nM]: 0 19 38 50 75 100 125 150 175 225 300
Fraction of uxuB bound

**Supplementary Figure S2. Mutational analysis for the *uxuB* leader sequence by EMSA.** The secondary structure of uxuB was predicted using the Vienna RNA webserver. Mutations were designed to preserve the secondary structure and base-pairing probability of the overall structure. Binding sites considered for analysis (yellow) and the introduced mutations (red) are shown next to the structure.

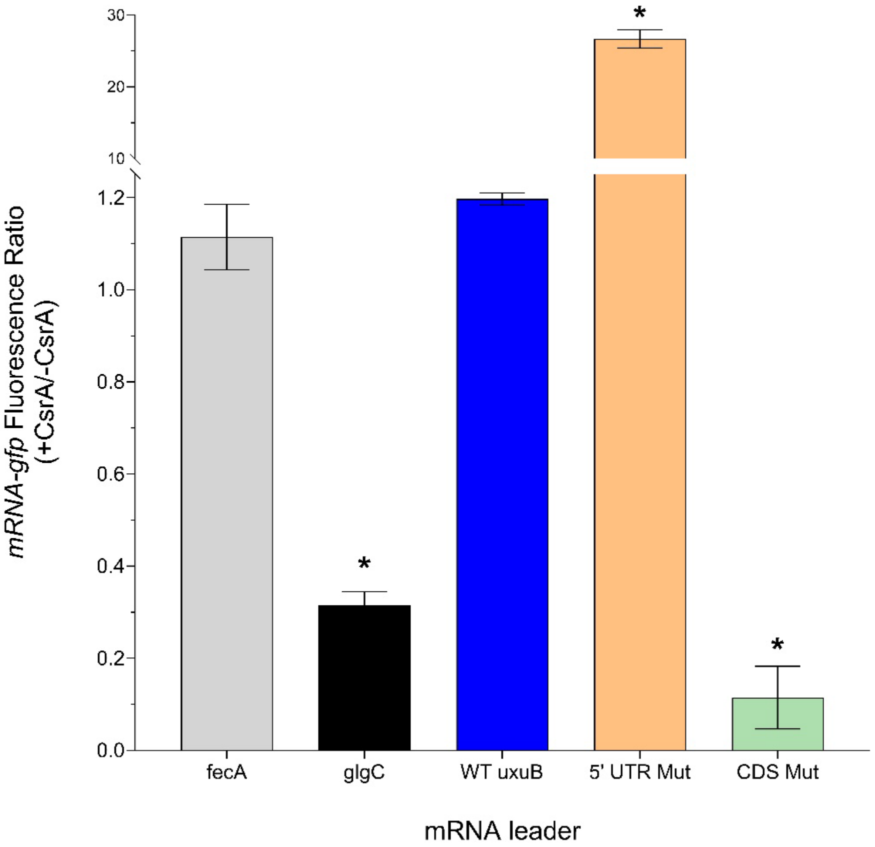

**Supplementary Figure S3. *In vivo* reporter assay for 5’ UTR and CDS Mutant *uxuB* leader sequences.** The 5’ UTR Mutant and the CDS mutant *uxuB* leader sequences were fused to *gfpmut3* and constitutively expressed from a plasmid. Cultures were then either induced with IPTG to express CsrA from another plasmid, or not induced and fluorescent signal was measured 3 hours after induction. To evaluate regulatory impact, the fluorescence ratios were taken by dividing fluorescence of the cultures that expressed CsrA and the *uxuB* mutant-*gfpmut3* fusion by the culture that only had the *uxuB* mutant-*gfpmut3* fusion. A ratio less than one indicates translational repression by CsrA, a ratio greater than one indicates translational activation by CsrA. Asterisks designate statistical significance in fluorescent ratios compared to the negative control, fecA, using a heteroscedastic t-test, in which ($p$-val $< 0.05$).

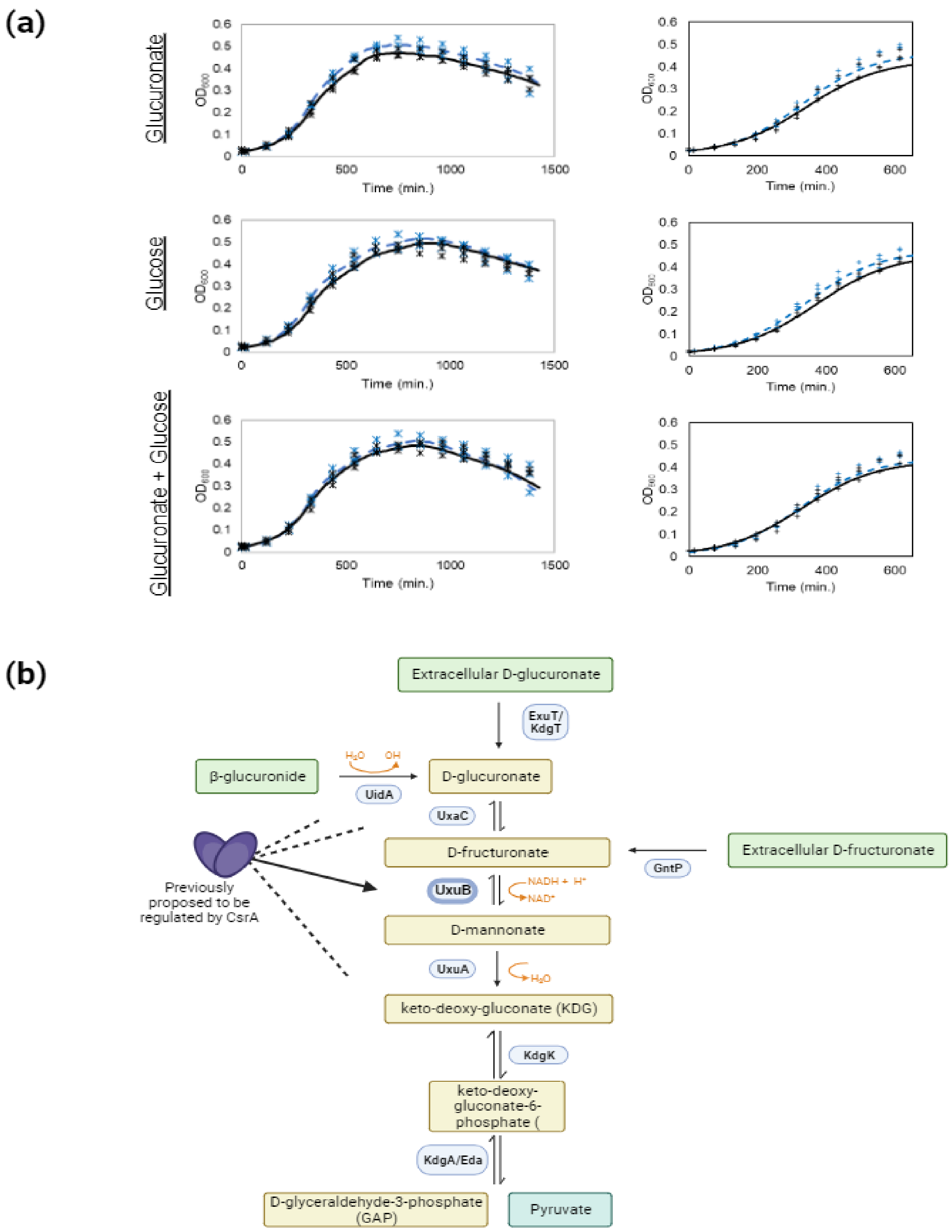

**Supplementary Figure 4. The CsrA-*uxuB* interaction does not significantly impact utilization of hexuronic acids.** (a) Testing for changes in the diauxic shift in the wild type vs the no GGAs mutant during growth on glucuronate (top), glucose (middle), or both sugars (bottom) The lack of shift from wild type to mutant suggests no change in regulation. (b) Proposed integration of CsrA regulation within the D-glucuronate catabolic pathway. Figure created with Biorender.com.

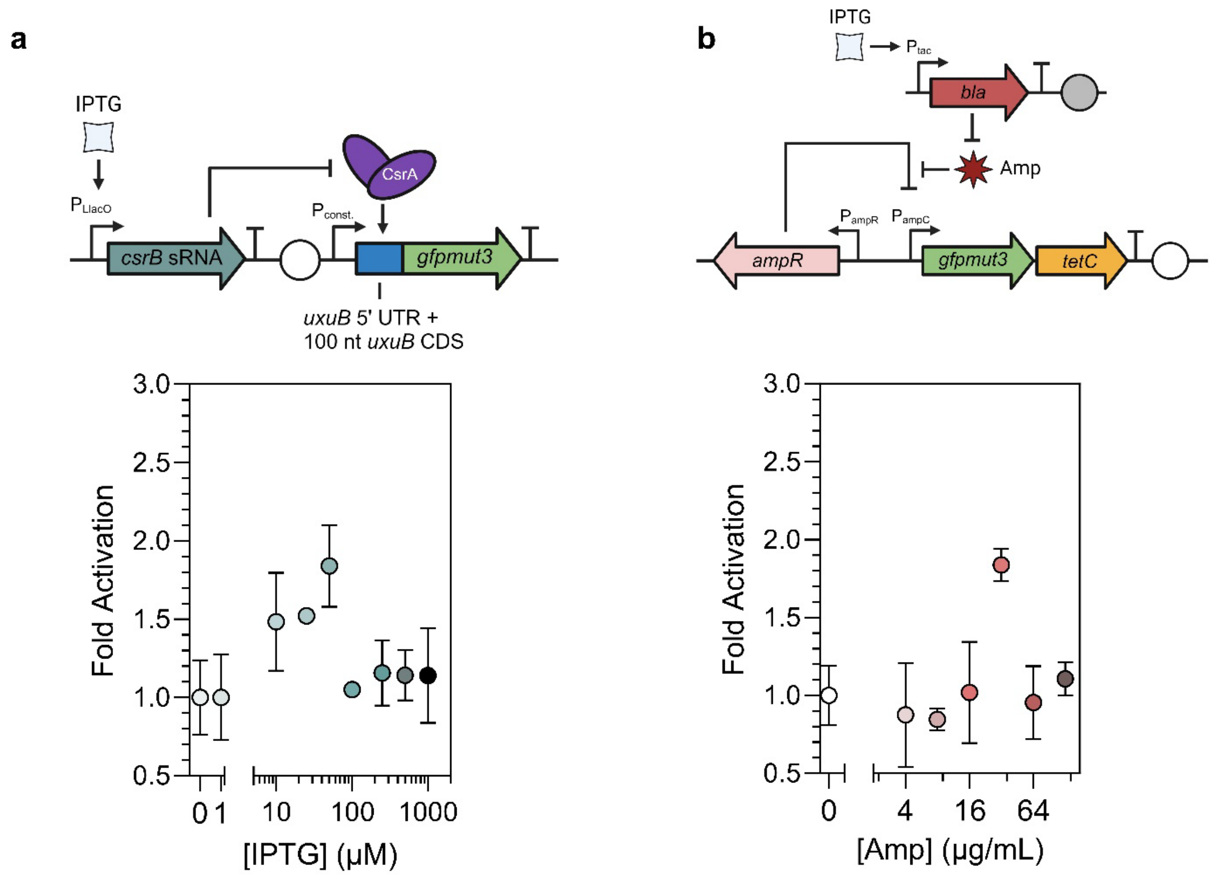

**Supplementary Figure S5. Benchmarking the Csr-regulated BANDPASS Filter against a transcriptionally regulated BANDPASS filter.** a) Genetic diagram of the Csr-regulated BANDPASS Filter and fold activation across an IPTG titration. b) Genetic diagram of the transcriptionally regulated BANDPASS filter developed by Sohka and colleagues[8] and the fold activation of the system using an Ampicillin titration.

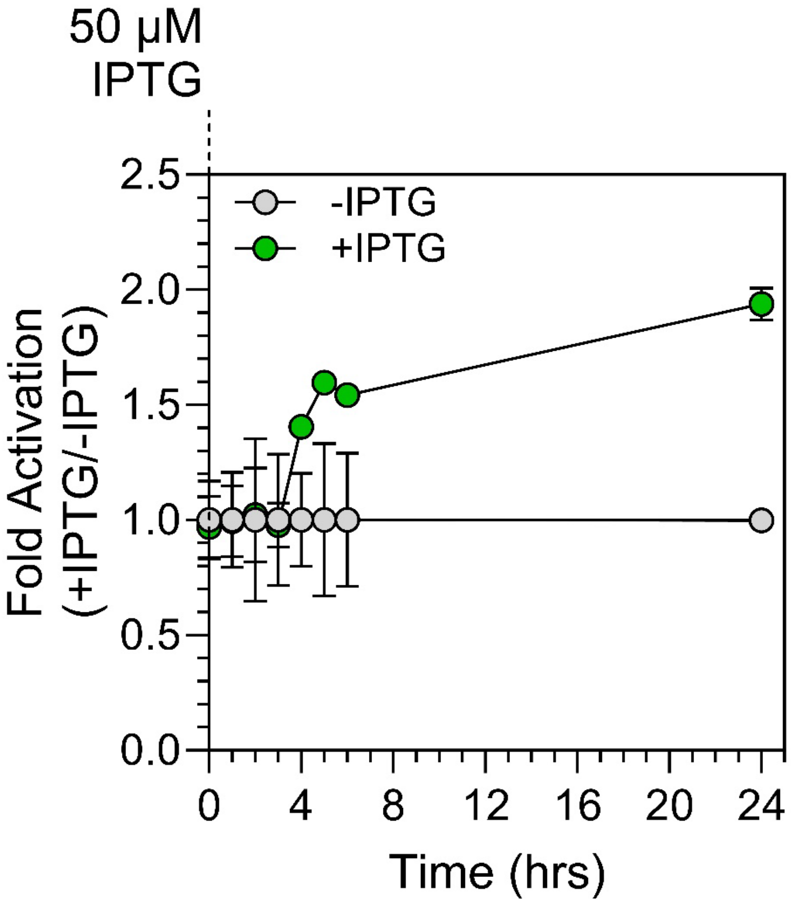

**Supplementary Figure S6. Time course of BANDPASS Filter fold activation at different induction levels.** Samples were either induced at seeding using 50 µM IPTG or uninduced.

**Supplementary References.**
